# Cognitive and non-cognitive outcomes associated with student engagement in a novel brain chemoarchitecture mapping course-based undergraduate research experience

**DOI:** 10.1101/768465

**Authors:** Christina E. D’Arcy, Anais Martinez, Arshad M. Khan, Jeffrey T. Olimpo

## Abstract

Course-based undergraduate research experiences (CUREs) engage emerging scholars in the authentic process of scientific discovery, and foster their development of content knowledge, motivation, and persistence in the science, technology, engineering, and mathematics (STEM) disciplines. Importantly, authentic research courses simultaneously offer investigators unique access to an extended population of students who receive education and mentoring in conducting scientifically relevant investigations and who are thus able to contribute effort toward big-data projects. While this paradigm benefits fields in neuroscience, such as atlas-based brain mapping of nerve cells at the tissue level, there are few documented cases of such laboratory courses offered in the domain.

Here, we describe a curriculum designed to address this deficit, evaluate the scientific merit of novel student-produced brainatlasmapsofimmunohistochemically-identifiednervecellpopulations for the rat brain, and assess shifts in science identity, attitudes, and science communication skills of students engaged in the introductory-level Brain Mapping and Connectomics (BM&C) CURE. BM&C students reported gains in research and science process skills following participation in the course. Furthermore, BM&C students experienced a greater sense of science identity, including a greater likelihood to discuss course activities with non-class members compared to their non-CURE counterparts. Importantly, evaluation of student-generated brain atlas maps indicated that the course enabled students to produce scientifically valid products and make new discoveries to advance the field of neuroanatomy. Together, these findings support the efficacy of the BM&C course in addressing the relatively esoteric demands of chemoarchitectural brain mapping.

## 1. Introduction

Course-based undergraduate research experiences (CUREs) are broadly recognized as an effective active-learning approach in a laboratory context (Chickering & Gamson, 1987; Bonwell & Eison, 1991; National Research Council, 1996; 2003; Handelsman et al., 2004; American Association for the Advancement of Science, 2011; PCAST, 2012; Auchincloss et al., 2014). Several key aspects distinguish CUREs from other active-learning approaches and undergraduate research experiences. Briefly, through a collaborative and iterative process, CUREs engage students in scientific practices that promote discovery of novel findings that are of broader relevance to the scientific community (Auchincloss et al., 2014). The resulting shifts in course enjoyment, research skills development, autonomy, and retention observed for CURE students versus their non-CURE counterparts have motivated interest in developing CURE curricula across the STEM fields (Badalà et al., 2013; Auchincloss et al., 2014; Jeffery et al., 2016; Olimpo et al., 2016; Rodenbusch et al., 2016; Ballen et al., 2017; Dolan, 2017; Frantz et al., 2017). In neuroscience, while problem- and research-based courses within the subdisciplines of medicine (medical neuroanatomy), psychology, and cell/molecular biology are represented in the literature, a brief review of published curricula indicates a gap in CURE offerings designed for entry-level students (please refer to **Supplemental Materials I** for a description of such approaches). Moreover, none of the curricula reviewed approached big-data problems in neuroscience, nor engaged students in atlas-based neuroanatomical mapping. The limited presence of such CUREs may be attributable to a prevailing assumption that such nuanced work requires extensive knowledge of neuroanatomy that can only be mastered through upper-division coursework. Yet, if lower-division CUREs could be designed to successfully incorporate laboratory-based curricula that involve students acquiring basic atlas-based neuroanatomical mapping expertise, the advantages afforded to the research community by effectively crowd-training this labor-intensive process would be significant.

We deliberately use the term “crowd-training” instead of “crowdsourcing” to acknowledge that a training component is required before individuals could perform the specific type of neuroanatomical mapping techniques we describe here. Given that crowd-trained individuals (students in CUREs) have helped advance big-data projects in the field of genome annotation (Chen et al., 2005; Call et al., 2007; Elgin et al., 2016), and that certain projects within the sub-domains of pathology (Della Mea et al., 2014) and neuroanatomy (Roskams & Popović, 2016; Irshad et al., 2017; also see Oleson, 2011) have been crowdsourced successfully outside of a formal CURE environment, we reasoned that it was possible to develop, implement, and evaluate a CURE that provided students the means to conduct authentic scientific research in atlas-based neuroanatomical mapping.

Accordingly, with the support of a grant from the Howard Hughes Medical Institute (HHMI) (Grant 52008125), we created an introductory biology CURE called *Brain Mapping & Connectomics (BM&C)*, the first CURE of its kind in the country. Recently concluding its fifth year (Martinez et al., 2019), the research goals for students in BM&C are to 1) use histological methods to label and identify the chemical phenotypes (‘chemoarchitecture’) of hypothalamic neurons in brain tissue; and 2) to generate high-spatial-resolution chemoarchitectural maps of these neurons and their axonal projections using a standardized reference atlas of the brain (Swanson, 2004). Through this process, the students make discoveries concerning novel patterns of distribution and relationships among neuronal phenotypes. In some instances, because a neuronal phenotype is exclusively found in only a single region of the brain, the patterns also include subsets of neural connections from cells of known origin, thereby contributing to the newly emerging discipline of connectomics. Coupled with future plans to incorporate more classical connectomics-related analyses, such as the mapping of neuroanatomically traced projections, we opted to title the course ‘*Brain Mapping & Connectomics’* to encompass these current and future aspects of the neuroanatomical analyses conducted in the course. Importantly, the patterns of chemoarchitecture analyzed by the students are being described for the first time in a documented spatial model of the brain, providing a foundation for future novel inquiry.

In this article, we present the curricular framework, pedagogical considerations, student affective outcomes, and student product assessment from three cohorts of the BM&C course. Where appropriate, we compare outcomes between students enrolled in the BM&C CURE and a non-CURE matched comparison group. Portions of these data have been presented in preliminary form (D’Arcy et al., 2016a,b). A report that focuses on the primary neuroanatomical data generated by multiple cohorts of BM&C students is being prepared separately based on our preliminary reports focusing on the neuroscience aspects of the project (Wells et al., 2015a,b; D’Arcy et al., 2016c; Flores-Robles et al., 2017; Burnett et al., 2018; Martinez et al., 2018; 2019).

### 1.1: Course requirements

BM&C is an equivalent substitution for the two-semester introductory biology laboratory course sequence. Pre-/co-requisites for the course include the general biology lecture series (BIO1: General Biology and BIO2: Organismal Biology) and a research fundamentals course designed to familiarize students with scientific literature and the inquiry process. It should be noted that, unlike the traditional laboratory courses that meet once per week with no option for extended hours, BM&C meets in two, three-hour sessions per week with additional optional hours offered on an as-needed basis depending upon room and instructor availability.

### 1.2: Course preparation

In an effort to ensure effective use of class time and to reduce histological error associated with the research process (Simmons & Swanson, 2009), we performed the following pre-course preparations:

a. Adult male Sprague-Dawley rats of consistent body weight at time of sacrifice were used to reduce size variation and to ensure that the sizes of the brains to be studied were similar to that used to produce the Swanson (2004) rat brain atlas.
b. Standardized fixation protocols and chemicals were implemented by A. M.
c. Sections were prepared by A. M. using consistent microtome blade settings (angle and thickness) and a consistent approach to plane-of-section adjustments.
d. The tissue blocking method, tissue section thickness, and the tissue collection scheme were standardized to provide students with well-curated series of experimental tissue.

### 1.3: Mapping system

As described previously (Khan, 2013; Khan et al., 2018a,b), a major limiting step in the unification of certain kinds of neuroscientific data has been the (largely unintentional) neglect of the neuroscience community to adopt a common spatial framework within which to integrate diverse datasets. This is especially true for datasets in the laboratory rat, a model that has been a mainstay of neuroscience research for at least the better part of a century (*e.g.*, Herrick, 1926; also see Table 4 in Khan, 2013). Indeed, numerous studies have been published on the expression patterns of key macromolecules within the rat brain, but few of these patterns have been mapped to a standardized brain atlas. This type of mapping for the distributions of macromolecules in the brain allows scientists to contextualize their findings in relation to other datasets mapped to the same reference space, thereby unleashing the predictive potential of chemoarchitectural studies (Khan, 2013; Khan et al., 2018a). Here, we opted to use the Swanson rat brain atlas (2004) as our spatial framework and to train students to perform key experimental and analytical procedures to identify, localize, and map expression patterns of key neuropeptides in the rat brain. We selected the Swanson atlas for several reasons: (1) a few studies, including those conducted by our own multi-institutional collaborative teams, have already been published that have utilized this spatial framework for mapping molecular expression patterns (Swanson et al., 2005; Yao et al., 2005; Geerling & Loewy, 2006; Kerman et al., 2007; Hahn, 2010; Zséli et al., 2016; Santarelli et al., 2018); (2) we have published the first explicitly-documented plane-of-section analysis for activity-dependent gene expression patterns using this atlas (Zséli et al., 2016), an analysis that is taught in this course; and (3) several studies are now taking advantage of the structured nomenclature system for this reference atlas, which can serve as an ontology for use by neuroinformatics specialists and computational neuroscientists seeking to integrate diverse datasets in digital space (see introductory comments in Brown & Swanson, 2013).

### 1.4: Course outcomes and learning/research objectives

We aligned course objectives to the aforementioned CURE dimensions (collaboration, iteration, scientific practices, discovery, and broader relevance), established CURE-specific outcomes to be assessed (research attitudes, science communication, science identity, and science process skills), and identified validated published instruments to be used in evaluating the specified affective outcomes (**Fig. 1, 2**) (Fredickson & Branigan, 2005; Estrada et al., 2011; Badalà et al., 2013; Auchincloss et al., 2014; Corwin et al., 2015; Hanauer and Hatfull, 2015; Weston and Laursen, 2015; Rodenbusch et al., 2016). We established clearly-defined, attainable minimum research objectives for the course: 1) to distinguish novel and distinct neuron populations in the rat hypothalamus by their chemoarchitectural properties; and 2) to represent the immunohistochemically-identified cells and fiber distributions in a canonical atlas for at least one complete atlas level. To aid in maintaining consistency across multiple cohorts taught by four different instructors, we developed a scoring rubric for student-generated maps. We also identified a suite of technical skills specific to the course research project as well as broadly-applicable research skills including documentation, science literacy, and science communication; all necessary to achieve project success.

**Figure 1.**
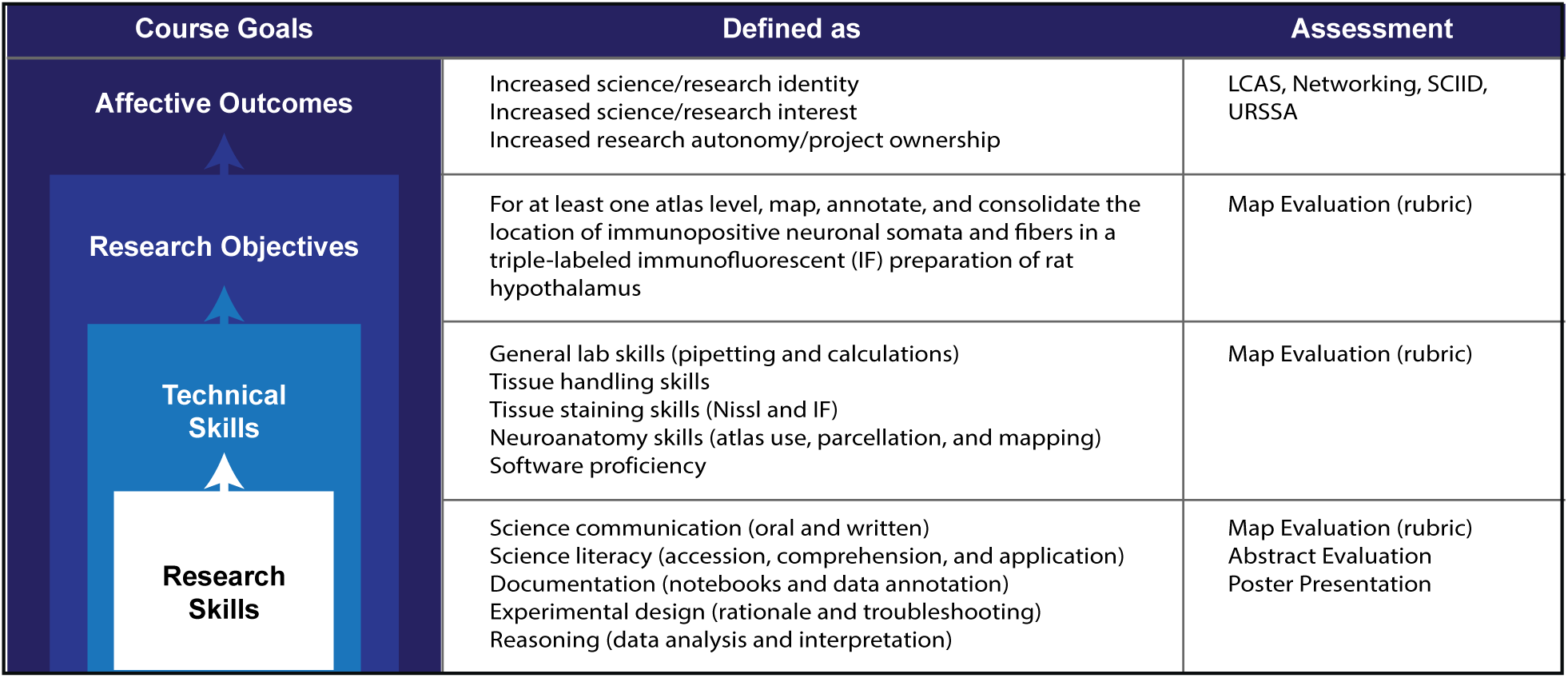
Course objectives and outcomes. By enabling students to practice research skills to gain self-sufficiency, to gain proficiency in a specific set of technical skills through iteration, and by setting attainable research goals, the course objectives were designed to yield positive student affective outcomes. Specific benchmarks were identified to describe successful outcomes for each tier of objectives. The mechanism employed to assess each outcome is provided. Laboratory Course Assessment Survey = LCAS; Science Identity Scale = SCIID; Undergraduate Research Student Self-Assessment = URSSA.

**Figure 2.**
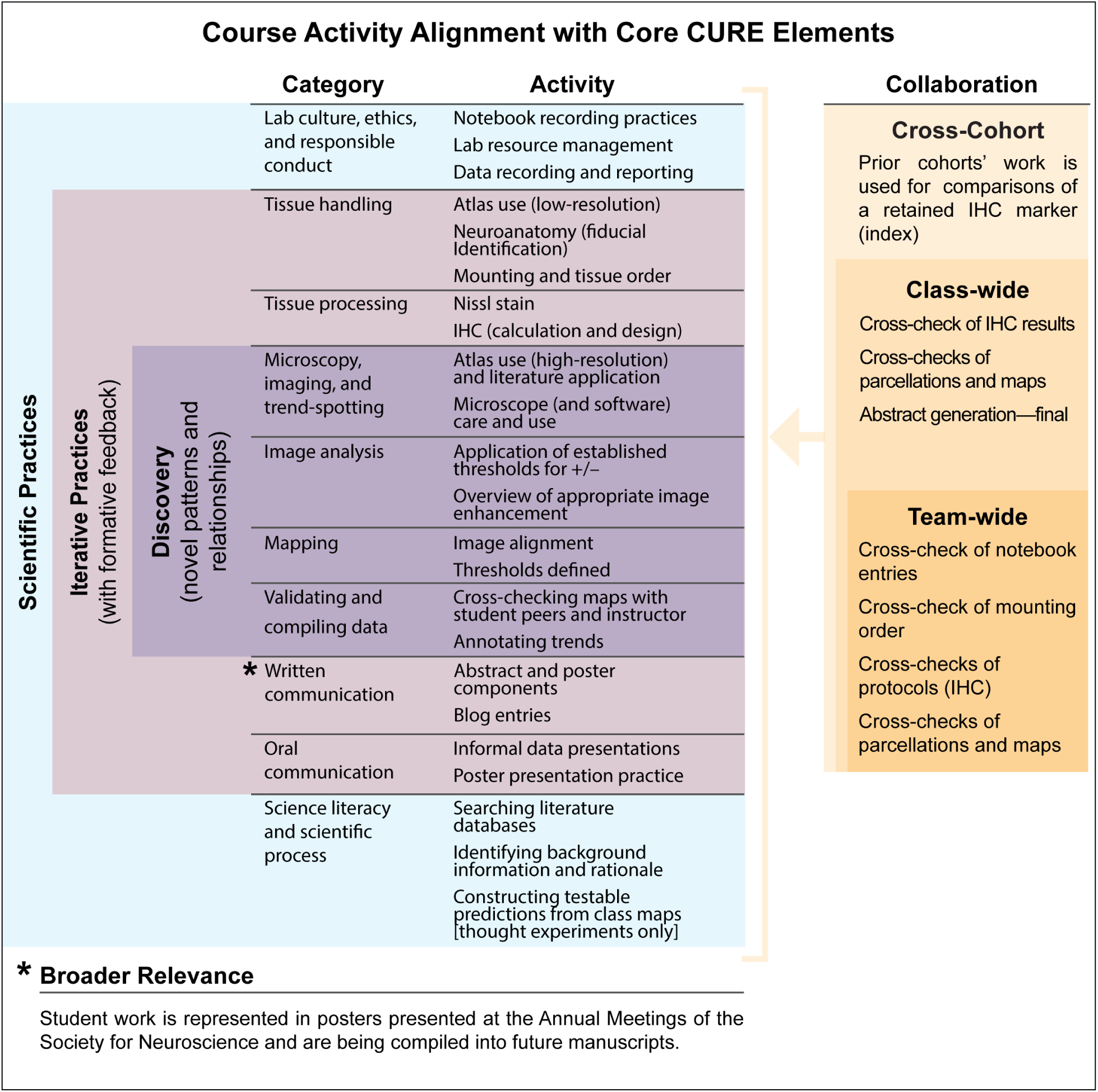
Alignment of course activities to the five dimensions of CUREs. Course activities were aligned to the five classic CURE elements: Scientific practices, iterative practices, discovery, collaboration, and broader relevance. Specifically, students iteratively engaged in authentic scientific practices within a tiered framework of collaboration. Their goal was to identify and contextualize novel chemoarchitectural patterns of neurons within hypothalamic regions of interest.

### 1.5: Course description and implementation

#### 1.5.1: Establishing student autonomy

Pedagogy progressively shifted from an instructor-driven style in the first weeks of the course (roughly four sessions in each cohort), to the demonstration and application of techniques, and finally to peer training. The peer-training model capitalized upon the staggered progress of individual groups. Teams that completed their protocols early were given instruction in new techniques, which they could then share with other teams, thus providing a means by which focused mentorship could be effectively scaled up while promoting peer teaching and actively demonstrating time-management skills. Finally, students moved toward class-wide collaboration and provision of feedback on generated products (*e.g.*, maps).

#### 1.5.2: Establishing research skills

Students were placed in permanent groups (n_members_ = 4) at the beginning of the course and provided with a general introduction to neuroanatomical terminology, the course goals, lab safety, and expectations of how a laboratory notebook should be used (**Fig. 2**). Instructors candidly discussed the tissue sample origins and introduced the policies of humane animal handling as well as the IACUC standards for animal care and use. Instructors also informed students that, while care should always be taken in carrying out lab protocols, mistakes inevitably occur. Students were encouraged to treat the errors they made as an important part of their learning process and not something to fear or to obfuscate. In an effort to reinforce this mentality, students participated in developing a strategy for error reporting, recording, and evaluation. Each class session began with a student-driven review of the key points of the preceding lab session, as recorded in their notebooks, and an overview of the current class objectives. Each class session ended with an instructor-led preview of the next set of planned class activities. Instructors used the time during tissue incubation periods to introduce new skills, assign students small projects (thought experiments, abstract and poster development activities, or informal in-class presentations on principles and techniques), and to share interesting news and information relevant to course content.

#### 1.5.3: Immunohistochemistry and tissue processing

Students practiced skills using non-critical tissue until they reached the desired proficiency level (two rounds practicing the mounting of tissue sections onto microscope slides and two rounds practicing immunohistochemistry provided students with sufficient training and troubleshooting opportunities to proceed on to using critical tissue). Once student proficiency was sufficient to reduce intra-group variability, instructors assigned an experimental rat brain – pre-cut into 30 µm-thick coronal-plane sections – to each group for further processing. Each team within a cohort performed multi-label immunohistochemistry of rat brain tissue sections for an identical set of three distinct antigens (neuropeptide biomarkers; **Table 1)**. This approach afforded students the ability to troubleshoot techniques together as a class, draw comparisons of immunopositive staining across four different animals, and identify robust staining patterns during class-wide examination of preliminary images. Students then worked to image tissue sections in the region bounded by the rostrocaudal extent of the lateral hypothalamic area stitched at ×10 magnification using a Zeiss Axio Imager M.2 epifluorescnce microscope (Carl Zeiss Microscopy, LLC; Thornwood, NY) driven by Neurolucida software (MBF Bioscience; Williston, VT). Students matched immunofluorescence images with images of the adjacent Nissl-stained tissue section for use in parcellation and mapping, as described below.

**Table 1.** Primary antibodies by cohort*.

| Cohort | Functional association | Target & Host/Type | Antibody registry | Source | Cat # | Dilution |
| --- | --- | --- | --- | --- | --- | --- |
| 1,3 | feeding, arousal, metabolism | melanin-concentrating hormone (MCH); rabbit polyclonal | AB_10013632 | Phoenix Pharmaceuticals | H-070-47 | 1:20K |
| 1,4 | responsivity/signalling regulation | NOS1, mouse monoclonal | AB_626757 | Santa Cruz | sc-5302 | 1:5K |
| 1,2,5 | feeding, arousal, metabolism | alpha-melanocyte stimulating (aMSH), sheep polyclonal | AB_91683 | EMD-Millipore | AB5087 | 1:10K |
| 2,5 | responsivity/signalling regulation | calbindin (CB-955); mouse monoclonal | AB_1140073 | Abcam | AB66185 | 1:1K |
| 2,3,4,5 | feeding, arousal, metabolism | orexin A (C-19), goat polyclonal | AB_653610 | Santa Cruz | sc-8070 | 1:4K |
| 3 | reward | tyrosine hydroxylase, mouse monoclonal | AB_2201528 | EMD-Millipore | MAB318 | 1:2K |
| 4 | feeding, arousal, metabolism | Agouti-related Peptide (AgRP) [82–132]-NH<sub>2</sub>; rabbit polyclonal | AB_2313908 | Phoenix Pharmaceuticals | H-003-53 | 1:2K |

\* Students retained at least one common antibody to act as an index for between-cohort variability in staining, imaging, and subject.

#### 1.5.4: Neuroanatomical mapping

Instructors introduced Adobe Photoshop CC and Adobe Illustrator CC (Adobe Systems, Inc., San Jose, CA) skills through a series of simple exercises and provided students with a general style template (layer organization and naming, line thickness and marker settings) to scaffold their mapping efforts. Prior to mapping, students were assigned seminal research articles pertaining to the immunohistochemical markers they probed, in relation to the region of interest being examined. This approach was used to not only inform students of the expected outcomes, but also to highlight novel findings as students progressively mapped the regions. This information was then used for the informal, in-class development of novel research questions and experimental design, as well as contextualization of student findings on final posters or oral presentations.

Students were also provided with digital files containing the Swanson (2004) reference atlas and neuroanatomical literature references for the regions of interest in the rat brain. Neuroanatomical boundaries and immunopositive signal (both axonal fibers and neuronal perikarya) were marked directly in Adobe Illustrator CC onto data layers that were superimposed over the layers containing photomicrographic images of immunofluorescence- and Nissl-stained tissue sections (**Fig. 2**). Students then transcribed all data onto the appropriate locations in the Swanson (2004) reference atlas, paying careful attention to preserve the relative distance, distribution, and orientation in the rectified image and to perform plane-of-section analysis. Finalized versions of all maps were assessed and validated by a rubric developed for this purpose (**Supplemental Materials II**). As a culminating product for the course, students worked collaboratively with their peers and the course instructor to incorporate mapping data into a conference poster for presentation at the Annual Meeting of the Society for Neuroscience (years 2016–2018).

#### 1.5.5: Assigning grades

Throughout the course, students were assigned the tasks of writing small blog entries, drafting abstract and poster content, and maintaining a laboratory notebook. They were also given occasional concept checks in the form of brief quizzes. Importantly, students were informed that the good-faith effort in producing quality research would be assigned a grade, not the project success. In this way, students were alleviated from the artificial pressure of grades that could potentially tempt them to behave dishonestly (Please refer to the course syllabus: **Supplemental Materials III**).

## 2. Materials & Methods

### 2.1: Student population

Students entered BM&C having met the pre-/co-requisite qualifications, as previously described. Three cohorts of BM&C students (n = 42; 79% Hispanic; 79% female) represented majors from multiple STEM disciplines, including: physics, psychology, forensic science, kinesiology, biochemistry, engineering (mechanical, biomedical, electrical), and biological sciences. Twenty-one percent of students were first-generation college attendees, and 50% reported English as their second language. From corresponding non-CURE biology laboratory courses, we drew a matched population for course comparisons, where indicated (n = 42). All participation was strictly voluntary, and surveys were conducted by non-instructor personnel. Students choosing to participate in surveys and data evaluation for the purpose of this study signed an informed consent document granting permission to use their responses in accordance with policies and procedures established by the university’s Institutional Review Board (IRB approval #789648). Unless otherwise indicated, all instruments were administered to participants at the start of the semester (pre-semester) and again at the end of the semester (post-semester) for pre *vs.* post comparisons.

### 2.2: Laboratory Course Assessment Survey (LCAS)

To gain a more complete inventory of the course environment, experience, and resources available to students, the LCAS was administered at the end of the course to both BM&C and non-CURE participants (Corwin et al., 2015). Responses to the 17-question survey were provided using a Likert-item scale (1= strong statement disagreement; 5 = strong statement agreement) and questions clustered into three categories (collaboration, discovery, and iteration) for the purposes of analysis. Psychometric data indicated a high level of construct validity and instrument reliability (Cronbach’s α ≥ 0.731 for all categories).

### 2.3: Networking (NW)

The Networking scale (Hanauer & Hatfull, 2015) was used to evaluate the extent to which BM&C and non-CURE students communicated about course content with individuals outside of the laboratory classroom environment. Student responses were recorded using a five-point Likert-item scale (1= strongly disagree; 5 = strongly agree). Students from both BM&C and non-CURE classes were surveyed at the end of their respective courses.

### 2.4: Science Identity Scale (SCIID)

We evaluated students’ science identity development using the Science Identity Scale employed by Estrada et al. (2011). The SCIID was selected intentionally given its previous use in evaluating historically-minoritized students’ science identity development in STEM contexts. Student responses were recorded on a five-point Likert-item scale (1 = strongly disagree; 5 = strongly agree). Surveys were administered to the BM&C and non-CURE cohorts in pre-/post-semester fashion. Psychometric analyses indicated a high level of instrument reliability (Cronbach’s α ≥ 0.837 for both pre- and post-assessment administration of the SCIID).

### 2.5: Undergraduate Research Student Self-Assessment (URSSA)

At the conclusion of the BM&C course, a modified version of the URSSA (Weston & Laursen, 2015) was administered to gauge students’perceptions of gains in four thematic dimensions related to scientific practices and professional development. These dimensions included: (a) thinking and working like a scientist [science process]; (b) personal gains; (c) research skills; and

(d) attitudes and behaviors [engagement]. Student responses were reported on a five-point Likert-item scale (1 = lowest or most negative rating; 5 = highest or most positive ranking). Scaling criteria were dependent upon the dimension being evaluated, as follows: (a) thinking and working like a scientist, personal gains, and skills (no gains - high gains); and (b) attitudes and behaviors (not at all - extremely frequently). Psychometric analyses indicated a high level of construct validity and instrument reliability (Cronbach’s α ≥ 0.848 for all dimensions).

### 2.6: Statistical analysis

Descriptive statistics were calculated for each dimension presented on the URSSA, with mean item scores reported. LCAS, NW, and SCIID responses were analyzed using a series of Analysis of Covariance (ANCOVA) and Multivariate Analysis of Covariance (MANCOVA) procedures, controlling for student demographics and pre-semester response (where appropriate). SPSS (v.23) was used to conduct all analyses.

### 2.7: Neuromapping Scoring Rubric (NSR)

Development of a rubric for the validation of student products was essential to: (1) examine student performance and proficiency with respect to brain mapping; (2) ensure scientific validity of student products; and (3) provide the graduate teaching assistant instructor with a standardized mechanism for scoring student products. Descriptions of each rubric scoring criterion are included in **Supplemental Materials II**. Briefly, rubric scoring fields are designed to account for varying levels of cognitive demand ranging from low-demand tasks such as attention to detail and following instruction (*e.g.*, using assigned formats of layer organization and naming conventions) to high-demand and highly-nuanced tasks (*e.g.*, correctly transcribing immunopositive label from the location in the experimental brains to the appropriate corresponding location in the atlas). Inter-rater reliability was assessed using de-identified student-generated maps that represented the hypothalamic region of Level 26 (Swanson, 2004) for two scorers (k = 0.88; *p* < 0.001). All disputes were resolved through iterative discussion between coders.

## 3: Results

In order to reduce potential bias associated with variation in laboratory graduate teaching assistant (GTA) instructor for both the BM&C and non-CURE course sections involved in this research, we first analyzed all outcome variables (*e.g.*, LCAS dimensions; science identity) using a Multiple Analysis of Covariance (MANCOVA) procedure. Results indicated no significant difference in student outcomes as a function of laboratory GTA instructor (F(42, 177) = 1.407; Wilks’ Λ= 0.259; *p* = 0.067; η_p_ ^2^ = 0.202). Furthermore, this effect was found to be true for both those GTAs facilitating the BM&C CURE (*p* = 0.573) as well as those GTAs facilitating non-CURE sections of the laboratory course (*p* = 0.199). Therefore, we aggregated data from all BM&C sections into a single cohort and likewise did the same for all non-CURE sections prior to conducting all subsequent analyses.

### 3.1: BM&C students report having greater opportunities to engage in discovery, collaboration, and iteration within the laboratory environment than their non-CURE peers

CUREs are designed to provide students with an opportunity to engage in novel research that is both collaborative and iterative in nature (Auchincloss et al., 2014). Consequently, we sought to understand the extent to which these design elements were present both within the BM&C CURE as well within the non-CURE learning environment. Students’ post-semester Laboratory Course Assessment Survey (LCAS) responses were analyzed using a series of ANCOVA procedures, controlling for student demographic attributes. Results indicated statistically significant, between-group differences on all outcomes: discovery (F(1, 76) = 13.710; *p* < 0.001), iteration (F(1, 78) = 6.340; *p* = 0.014), and collaboration (F(1, 72) = 31.680; *p* < 0.001); with those students enrolled in the BM&C CURE reporting greater opportunities to engage in the aforementioned processes than their non-CURE counterparts (**Fig. 3**).

**Figure 3.**
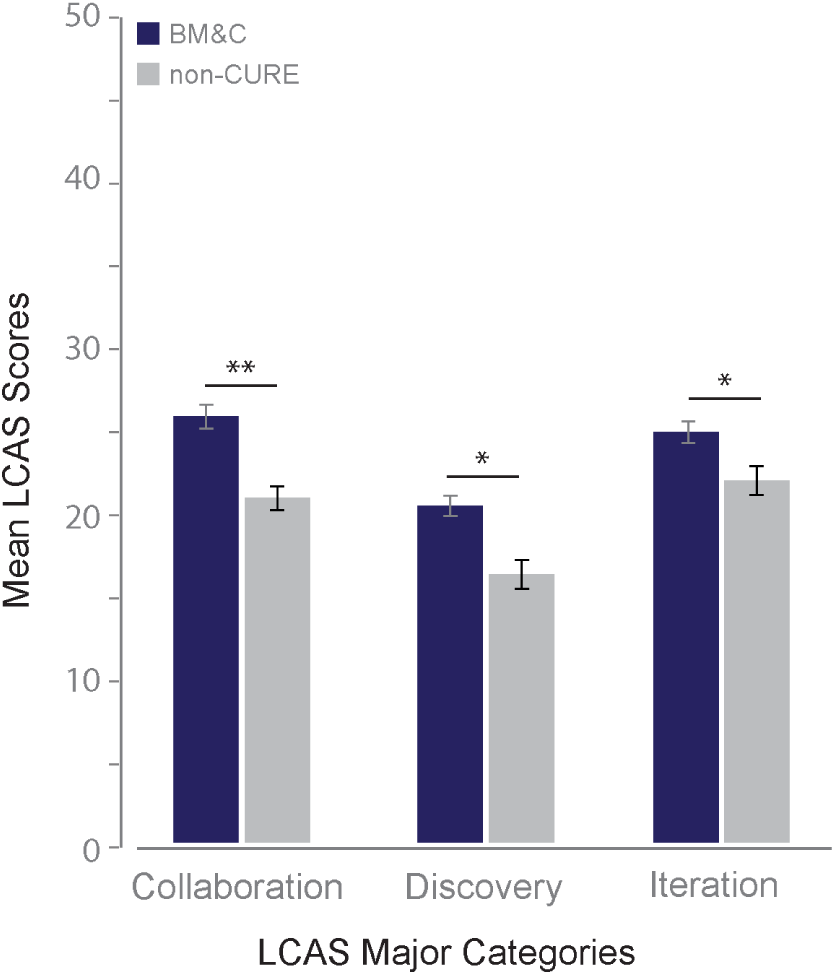
Students’ perceptions of the laboratory environment. In a matched comparison of traditional introductory biology laboratory courses (Non-CURE *in gray*) versus the BM&C (*in blue*) courses, BM&C students show significantly higher scores in collaboration (\*\**p* < 0.005), discovery and iteration categories (\**p* < 0.05).

### 3.2: BM&C students communicate about their research to external stakeholders more frequently than their non-CURE peers

Whilethe LCASprovidesavalidmeasureof within-course collaboration, we were also interested in exploring the degree to which BM&C and non-CURE students communicated information about their research to individuals external to their laboratory course environment. Post-semester data obtained from administration of the Networking Scale (Hanauer & Hatfull, 2015) were analyzed using a series of ANCOVA procedures, controlling for student demographic attributes.

The results (**Fig. 4**) revealed a statistically significant difference in composite score between the BM&C and non-CURE cohorts (F(1, 78) = 18.867; *p* < 0.001). Post-hoc MANCOVA analyses demonstrated further that BM&C students discussed their research with parents/guardians (F(1, 78) = 37.350; *p* < 0.001), friends (F(1, 78) = 20.733; *p* < 0.001), other students at their institution not participating in the BM&C course (F(1, 78) = 11.436; *p* = 0.001), and students at other institutions (F(1, 78) = 6.870; *p* = 0.011) to a greater extent than their non-CURE classmates.

**Figure 4.**
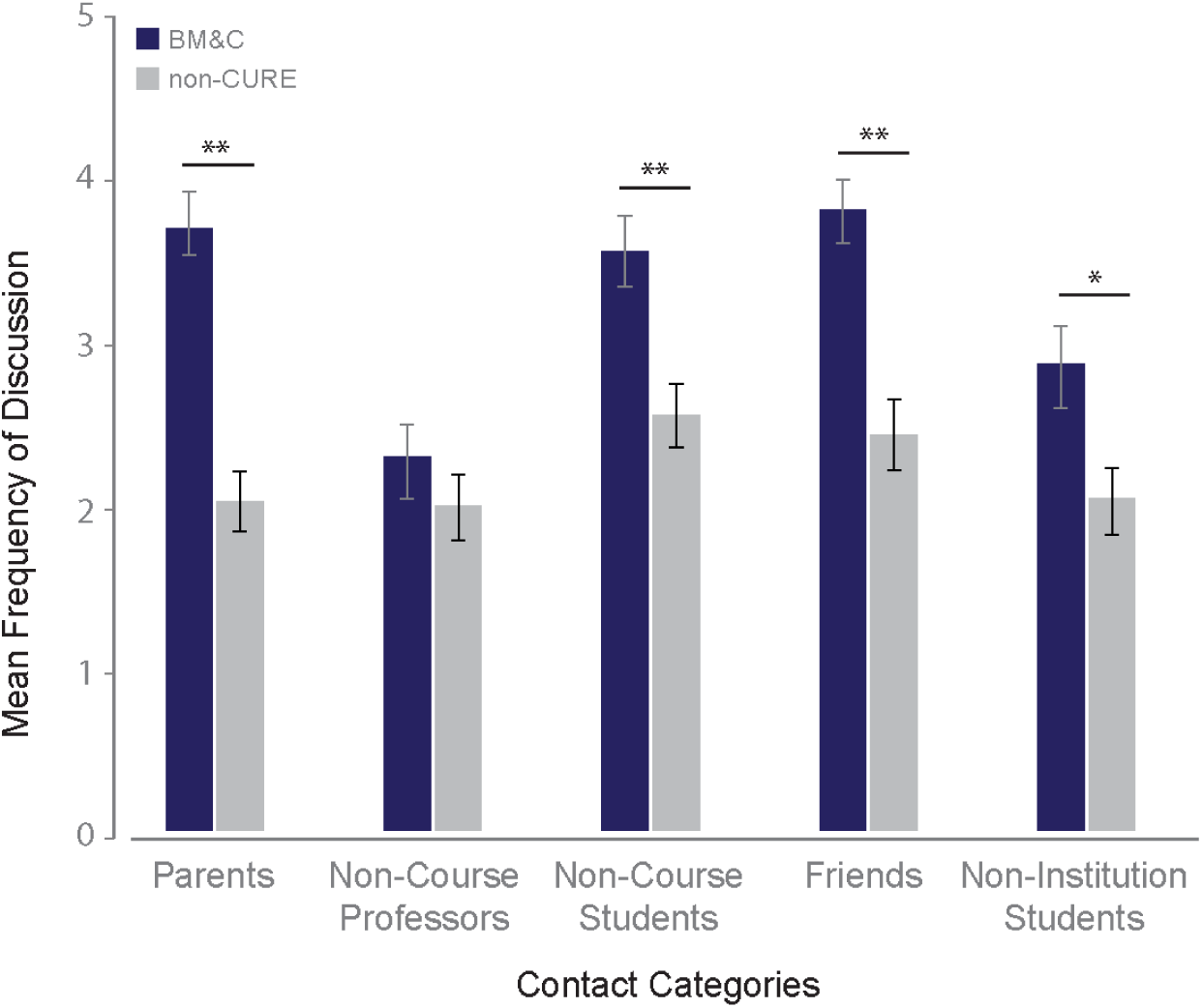
Student networking patterns. BM&C students (*blue*) engage in greater levels of external networking than their non-CURE (*gray*) peers (\*\**p* < 0.005).

### 3.3: BM&C students identify as scientists more strongly than their non-CURE counterparts

Given the role of science identity development as a positive mediator of student engagement, retention, and long-term persistence in STEM (Auchincloss et al., 2014; Estrada et al., 2011), we administered the Science Identity Scale (SCIID; Estrada et al., 2011) to participants in both the BM&C and non-CURE laboratory sections in pre-/ post-semester format. ANCOVA analyses, controlling for student demographic attributes and matched pre-semester SCIID response, demonstrated a significant, between-group difference in agreement on a single item: “I have come to think of myself as a scientist” (F(1, 77) = 10.177; *p* = 0.002); with BM&C participants reporting higher average statement agreement (**Fig. 5**).

**Figure 5.**
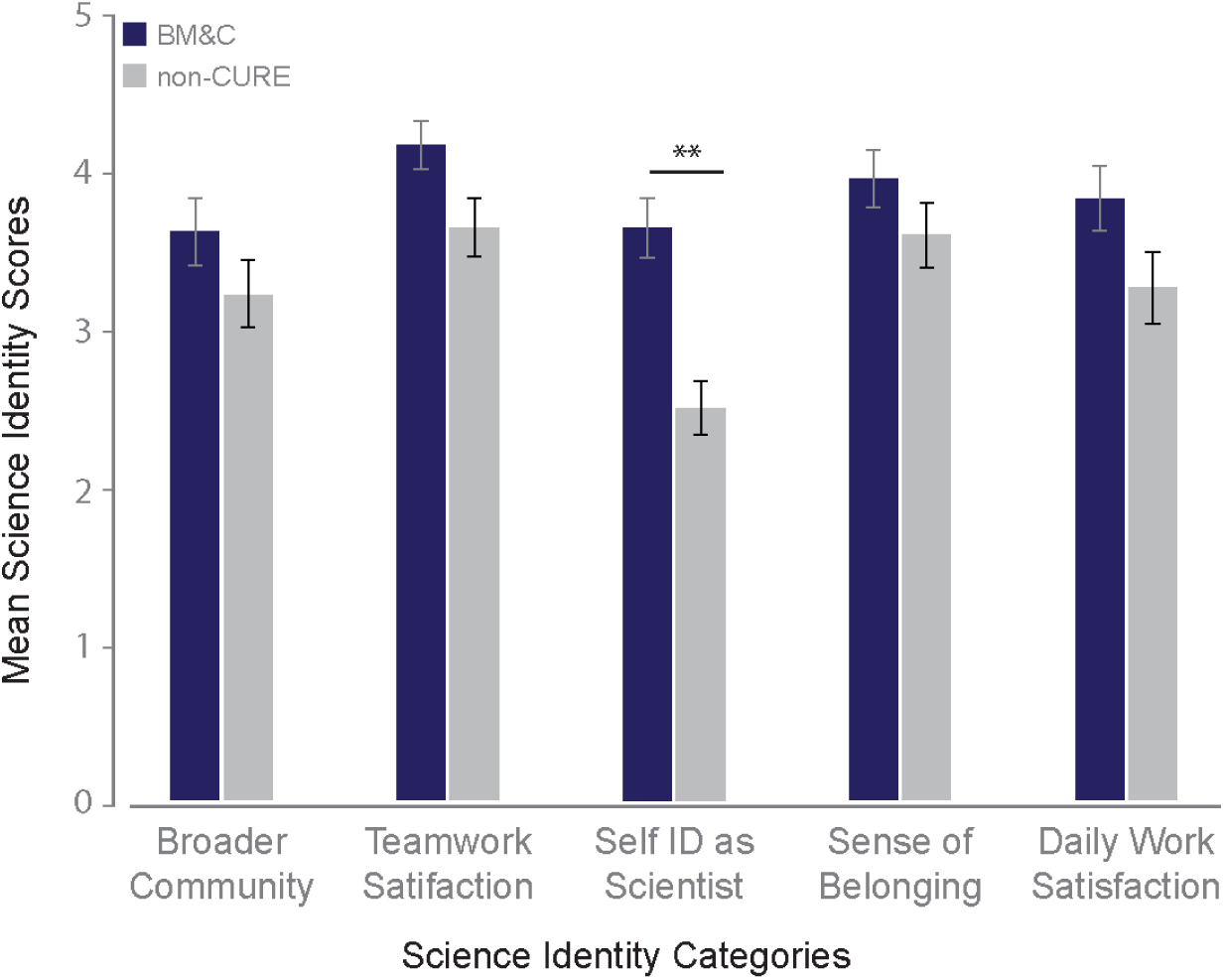
Science identity scores. While no differences were seen among many of the categories assessed, BM&C students (*blue*) did self-identify as a scientist more frequently than non-CURE students (*gray*) (\*\**p* < 0.005).

### 3.4: BM&C students demonstrate aptitude in atlas-based neuroanatomical mapping

Maps from each BM&C student team and iteratively evaluated using the Neuromapping Scoring Rubric (NSR) (an example of a completed rubric evaluation is provided in **Supplemental Materials II: Table SM2.1**; also see **Fig. 6A–C** for student workflow examples), which we developed specifically for this course. Students performed well in the *System and Logic* category, which evaluated their ability to follow the provided organizational guidelines (data layer order, naming, line and marker parameters). Students also generally exhibited good atlas and anatomical proficiency but required greater feedback and correction on plane of section analysis and boundary identification of certain sub-regions in the hypothalamus (*see* **Fig. SM2.3**). Prior to receiving instructor feedback, students likewise experienced difficulty correcting for tissue distortion when translating signal to the final map (*see* **Fig. SM2.3**).

**Figure 6.**
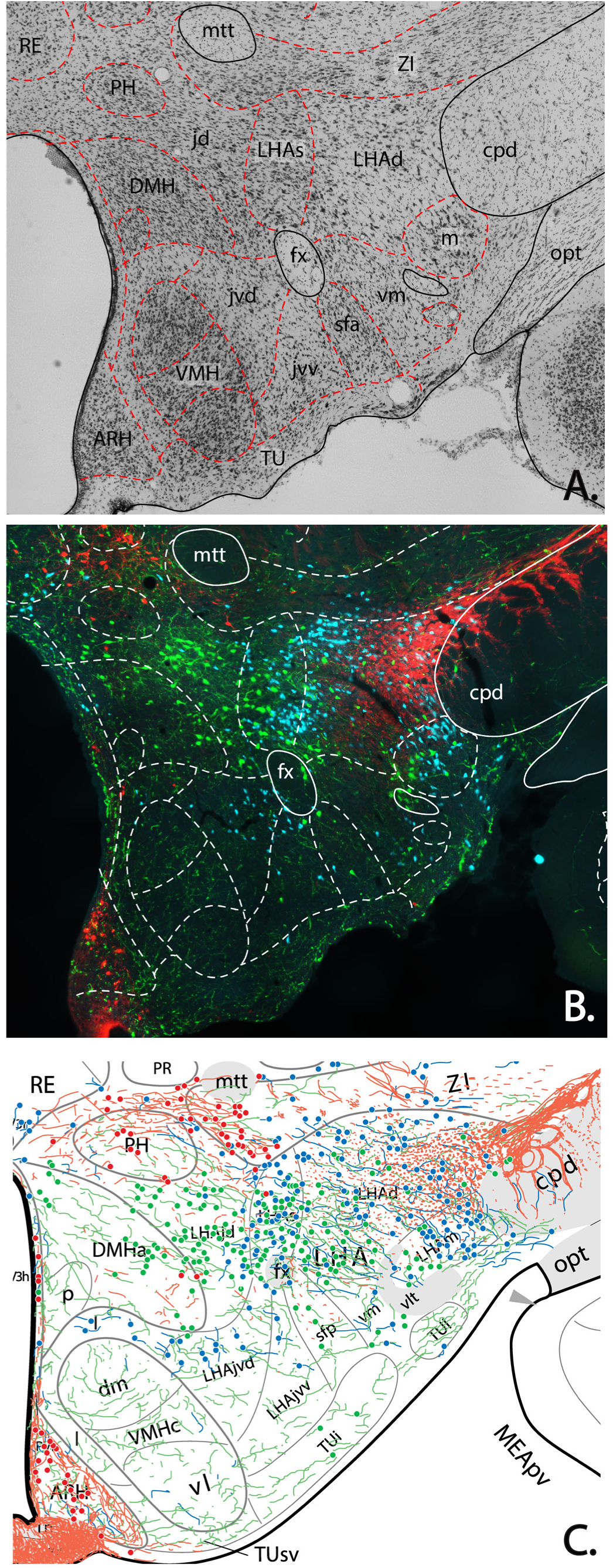
Sample of student workflow. **A**. Students identified structures based on the Nissl-stained cytoarchitecture (photographed at ×10 magnification). **B**. Boundaries from the Nissl-stained section were overlaid onto immunohistochemical data images (tyrosine hydroxylase (TH), *red*; melanin concentrating hormone (MCH), *blue*; hypocretin/orexin (H/O), *green*. All images taken at ×10 magnification). **C**. Merged data files of student-generated maps representing fiber density trends and denoting immunoreactive cells within the modeled framework. The underlying atlas template is from Swanson (2004) (available at https://larrywswanson.com) and is reproduced and modified here under the conditions of a Creative Commons BY-NC 4.0 license (https://creativecommons.org/licenses/bync/4.0/legalcode).

### 3.5: Student perceptions of the BM&C CURE

Post-semester Undergraduate Research Student Self-Assessment (URSSA; Weston & Laursen, 2015) data were obtained from individuals in the BM&C course in order to provide additional, nuanced detail about students’ experiences in the CURE and to provide formative feedback to the BM&C instructional team. Specifically, the URSSA prompts students to self-report gains in four areas: (a) Science Process; (b) Personal Gains; (c) Research Skills; and (d) Engagement. Descriptive analysis of mean scores within the *Science Process* category indicated that the greatest perceived gains were in finding patterns in data (*M* = 4.25; *SEM* = 0.12) and in being able to relate one’s research to his/her coursework (*M* = 4.53; *SEM* = 0.10) **(Fig. 7)**. A similar trend was observed for the *Personal Gains*-related questions, with BM&C students reporting good gains overall. Specifically, developing patience for the pace of research (*M* = 4.44; *SEM* = 0.12) and understanding what research is actually like (*M* = 4.52; *SEM* = 0.11) received the highest ratings. Subsequent analysis of student responses on *Research Skills* items demonstrated that BM&C undergraduates reported good gains for the majority of statements. Most notably, this included working with computers, conducting observations, maintaining laboratory notebooks, understanding journal articles, and explaining their project to people outside of their field. With respect to *Engagement*, participants reported that they frequently engaged in real-world research, felt responsible for their project, felt like a scientist, and thought creatively about their project (**Fig. 7**).

**Figure 7.**
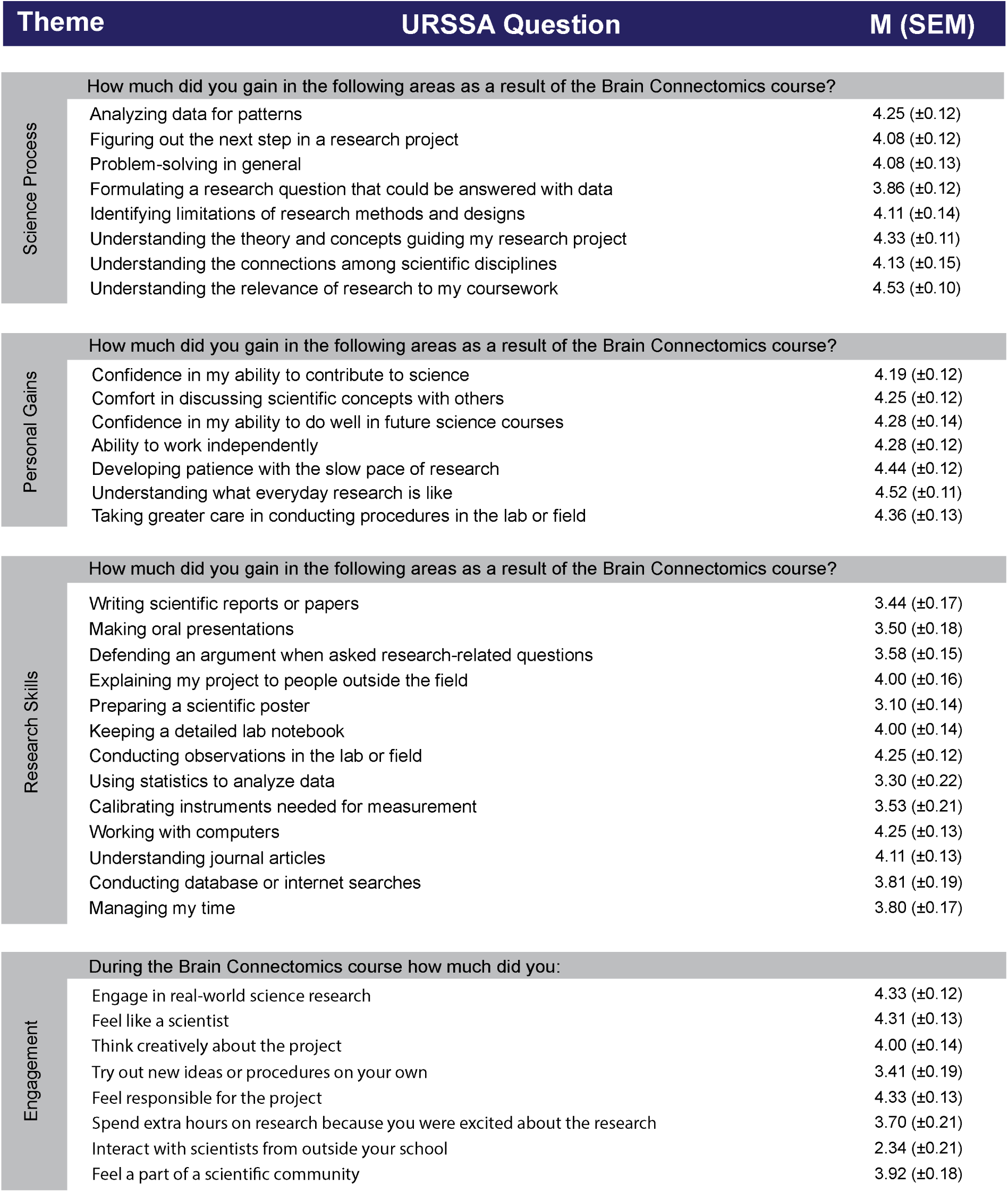
Post-semester URSSA responses. Questions are grouped by the theme assessed. M = Mean; SEM = Standard Error of the Mean.

## 4. Discussion

In this article, we demonstrate that the BM&C course, as implemented, provides students with many of the key positive affective outcomes associated with CUREs, including increased science identity, greater opportunity for advancement in technical skills, and increased research acculturation versus their non-CURE counterparts (Badalà et al., 2013; Hanauer and Hatfull, 2015; Jeffery et al., 2016; Olimpo et al., 2016; Ballen et al., 2017; Frantz et al., 2017). Moreover, these metrics are achieved in a crowd-training environment wherein students can perform high-quality research that can be readily disseminated in traditional circles of professional scientific communication (*e.g.*, scientific conferences; peer-reviewed publications). Indeed, some of the datasets produced by BM&C students have already been showcased at scientific conferences (Wells et al., 2015a,b; D’Arcy et al., 2016c; Flores-Robles et al., 2017; Burnett et al., 2018; Martinez et al., 2018). Together with the training rubric we have developed for evaluating atlas-based mapping of tissue labeling, this body of work constitutes an initial framework and proof-of-concept that can aid other investigators seeking to develop a full-length teaching laboratory centered upon atlas-based brain mapping at the mesoscopic (tissue-level) scale. Below, we discuss our results in the context of each of the evaluation and assessment mechanisms we employed in this study.

### 4.1: Impacts of the BM&C course on students’ science process skills and science identity development relative to non-CURE participants

The value of the authentic research experience is that it provides students with the ability to repeat laboratory work until they achieve a certain level of success or proficiency (iteration), grants students the freedom to pursue novel findings (discovery), and promotes greater interaction with team- and classmates in order to achieve progression in the project (collaboration) (Auchincloss et al., 2014; Corwin et al., 2015). These experiences can influence students’ willingness to speak to others outside the classroom about their research or shift the degree to which students perceive themselves to be a member of the scientific research community (Estrada et al., 2011; Hanauer & Hatfull, 2015).

Networking frequency of BM&C students was significantly greater among all peer and non-peer groups, excluding non-course professors, relative to their non-CURE counterparts (**Fig. 4**). This is consistent with Hanauer’s and Hatfull’s (2015) findings and suggests that BM&C students have personal interest and investment in the course activities to the extent that they commit time outside of class to think about and discuss their research with others. While outside of the scope of the present study, we contend that future research should examine the contextual factors mediating such networking behaviors.

With respect to science identity (Estrada et al., 2011), although BM&C and non-CURE students showed no significant differences in response across the majority of items, scores for both BM&C and non-CURE students were generally positive. Both class formats provided students with a moderate sense of belonging in science and within a broader scientific community. Likewise, students in both cohorts reported agreement with statements regarding teamwork and daily work satisfaction. Importantly, however, BM&C students did report a significantly higher propensity to think of themselves as scientists versus their non-CURE counterparts. Given the often-tedious nature of this form of neuroanatomical mapping, we were encouraged that BM&C students continued to find the daily work of a scientist appealing upon completion of the course.

### 4.2: Trends in student mapping performance

The Neuromapping Scoring Rubric (NSR) provided a mechanism by which BM&C instructors could score students’ initial mapping attempts and provide students with targeted formative feedback. From a pedagogical standpoint, implementation of the rubric also allowed instructors to identify aspects of the mapping process that posed a challenge for students and, thus, uncovered areas where future mentoring and feedback were required. For instance, one unanticipated observation was that students could easily identify immunopositive fibers with little need for correction but had intermittent success with cell body (neuronal perikaryon) identification. Often, students would mark cells that were *weakly immunopositive* (to be denoted with an open circle) as *immunopositive* (filled circle), thus indicating a need to create a class-wide guide for establishing thresholds. Much of the remaining correction required by students centered on drawing the anatomical boundaries of more subtly-defined sub-regions of the hypothalamus and on applying correction for tissue distortion when transferring data to the final maps (please refer to **Supplemental Materials II, Fig. SM2.2** and **Fig. SM2.3**, for further illustration of this application). In addressing anatomical boundary issues, instructors used class reference materials (in the form of additional examples of the regions of interest from their own work and from published literature) to highlight the key features of the cytoarchitecture that defined the regions. In transferring data to digital atlas templates, students required more extensive instruction in adjusting stretch and compression distortions before mastering the ability to correctly translate the spacing, relative positioning, and relative densities of signal (please see **Supplemental Materials II, Fig. SM2.4**, for an illustration of the described data transformation). However, students were able to act independently after one or two instructor-led interventions. Students were also observed providing increasingly meritorious peer critique after receiving instructor input, indicating increased concept mastery and process autonomy.

### 4.3: Student perception of gains experienced during the BM&C course

Students experienced moderate to good gains in many of the metrics assessed by the URSSA, with the highest gains observed in developing patience, conscientious research practices, and understanding the day-to-day realities of authentic research (*see* **Fig. 7**). These personal gains have been aligned with student acclimatization to the research environment (Weston & Laursen, 2015). Furthermore, in keeping with the course intent to empower students to engage in authentic research, students of the BM&C course report recognizing their work as “real-world” research, feeling like a scientist, and feeling responsible for their project.

### 4.4: Moving toward connectomics

As discussed in the Introduction, although the student research efforts catalogued here are limited in their relation to the “connectomics” portion of “Brain Mapping and Connectomics,” it was our deliberate choice to include this facet in the course title as we recognized the potential to expand the course into analyzing anterogradely- and retrogradely-traced rat brains to define the connections that exist between distinct neuron populations. In these initial cohorts, however, it was important to establish the proof of concept that novice students *could* perform and benefit from the course activities of immunohistochemically labeling, identifying, and accurately representing neuron and fiber populations within a modeled space. Furthermore, the student maps generated and vetted for these cohorts provide future students with the foundational data needed to begin hypothesizing potential networks among the neuron populations characterized to date within the literature. Using our tested curriculum framework, we are thus branching into tracing studies wherein students will discover and chart networks pertaining to feeding, memory, and fear (*e.g.*, Pineda Sanchez et al., 2019).

### 4.5: Scaling up BM&C

Although this study focuses on a small student population, group structures and the use of group members as peer-instructors can provide a means of scaling up class size while maintaining a single course instructor. In order to combine ever-larger sets of data across cohorts, we have developed a tool to assess student products, streamline evaluation and formative feedback, and unify clear expectations for the class. Multiple instructors, who have conducted BM&C courses in parallel to augment crowd-training and expedite data processing, have been aided greatly by the use of discussion forums and blogs to share feedback and upload systematic evaluation tools, appropriate brain mapping training materials, and a unified research protocol. Training workshops, in which instructors are equipped with the appropriate starting materials (*e.g.*, tissue sets) along with sample training materials, are recognized as a means to ensure consistency of performance longitudinally as has been done with next-generation sequencing projects (Buonaccorsi et al., 2011, 2014). Fostering intra- and extramural collaborations likewise offers a mechanism for leveraging the BM&C curriculum framework to advance student learning and success in resource-challenged classrooms. We discuss this potential in greater detail below.

### 4.6: Expansion, opportunity and accommodation through collaborations

Possible barriers to offering the BM&C course including animal subject availability, protocol approval and housing, available “wet lab” space, funding for antibodies and reagents, specific laboratory equipment (*e.g.*, microtome, fluorescence microscope); may be overcome through identifying collaborative exchanges within or among campuses. By way of example, pre-sectioned animal tissue can be provided by an external laboratory for CURE students to process (mount, stain, image, and map) requiring little modification to the curriculum as presented here. Alternatively, images of brain sections that have been stained and photographed by collaborating experts could be provided to CURE participants, thus allowing this curriculum to be developed as a “dry lab” that requires only computers, tablets, and Adobe software (Illustrator and Photoshop) for students to engage in the mapping-related activities. Thus, the bare minimum of equipment required by the course may vary depending on the specific resource gaps and the structure of the collaboration.

Finally, collaborations framed as described above can provide several unique dimensions to the CURE experience. Instructors can choose to incorporate professional networking and oral science communication activities in the form of online video lab meetings with collaborating students, staff, and faculty. Written communication products in the form of reports, well-curated notebooks, and data stewardship are likewise strongly emphasized under these circumstances. These approaches further promote a more inclusive experience as all participants have potential access to an expansive network of peers and experts who can effectively guide them in their research and broader professional efforts.

### 4.7: From crowd-trained research to crowd-directed research

The late physicist, Richard Feynman, remarked famously how students can often provide much insight into new research directions, despite having little or no expertise in the subject:

The questions of the students are often the source of new research. They often ask profound questions that I’ve thought about at times and then give up on, so to speak, for a while. It wouldn’t do me any harm to think about them again and see if I can go any further now. The students may not be able to see the thing that I want to answer, or the subtleties I want to think about, but they *remind* me of a problem by asking questions in the neighborhood of that problem. It’s not easy to remind yourself of these things. (Feynman, (1985), p 166).

From our experiences teaching BM&C, we found numerous occasions where inquisitive students would provide the types of “reminders” Feynman refers to, in which gaps of knowledge within the field are exposed during their discussions with instructors of the course. As alluded to previously, a promising area of expansion for our course would be to capture and harness the products of such discussions to generate new research questions and to have students help design and conduct new studies based on such questions.

In conclusion, with careful planning and realistic goals, novice undergraduates can generate quality high-spatial resolution chemoarchitectural maps of the brain. Importantly, our findings suggest that a neuroscience focus or major is not an essential requirement for students to derive satisfaction, research identity, and a sense of autonomy from conducting this type of research. Providing students with a sense of community, independence, and meaningful collaborative experiences can influence development of other affective and research-related outcomes.

## Acknowledgments

Evaluation surveys (URSSA and pre- and post-semester surveys) were developed and administered by the Research Evaluation & Assessments Service at the University of Texas at El Paso (UTEP), led by Dr. Guadalupe Corral. Care of experimental animals used in preparation for course materials was provided by the Laboratory Animal Resources Center at UTEP under a protocol approved for the UTEP Systems Neuroscience Laboratory by the UTEP Institutional Animal Care and Use Committee. We would like to thank Kenichiro Negishi for his assistance in validating the Neuromapping Scoring Rubric. We thank Drs. Joanne T. Ellzey, Elizabeth Walsh, Robert A. Kirken, Renato Aguilera, Craig Tweedie, and Kristine Garza for facilitating space allocation for the Brain Mapping & Connectomics laboratory; Drs. Stephen Aley and Bruce S. Cushing for supporting equipment procurement for the course; and Briana E. Pinales for work during the initial stages of the research project. We are also grateful to Dr. Larry W. Swanson for his thoughtful discussions and informal evaluations concerning the course, and for visiting with and providing feedback to our BM&C students at UTEP. The Brain Mapping & Connectomics CURE was created under the UTEP PERSIST (Program to Educate and Retain Students in STEM Tracks) program, funded by HHMI grant 52008125 (PI: S. Aley, Co-PIs: L. Echegoyen, A. M. Khan, D. Villagrán, E. Greenbaum). CED and AM were supported by the HHMI PERSIST program. Work in the UTEP Systems Neuroscience Laboratory was supported by NIH Grant GM109817 awarded to AMK; this grant also supported the procurement of supplies and equipment for the BM&C course. The UTEP Department of Biological Sciences and Office of Research and Sponsored Projects also supported equipment procurement for the course. Education research equipment, supplies, and personnel were supported, in part, by the BUILDing SCHOLARS program (NIGMS/NIH Award Numbers RL5GM118969, TL4GM118971, and UL1GM118970).

## D’Arcy *et al*. Supplemental Materials I: Literature Evaluation of CURE and Problem-based Curricula for Neuroscience Undergraduates

In searching extant literature describing neuroscience-themed CURES or inquiry-based approaches, the following search terms were applied in Google Scholar, ERIC, and NCBI’s PubMed database search engines: [“Undergraduate” AND “Research” AND “Neuroscience”] or [“Undergraduate” AND “Research” AND “Neuroanatomy”]. Articles were then assessed for content suitability, and courses specifically designed for medical neuroanatomy purposes, neuroscience program descriptions, novel distribution of teaching responsibilities, and non-inquiry-based courses were excluded. **Table SM1.1** (*below*) lists represents a non-exhaustive treatment of the literature, but provides a sense of focus and implementation in readily-available curriculum resources in neuroscience CUREs and problem-based courses.

**Table SM1.1:**
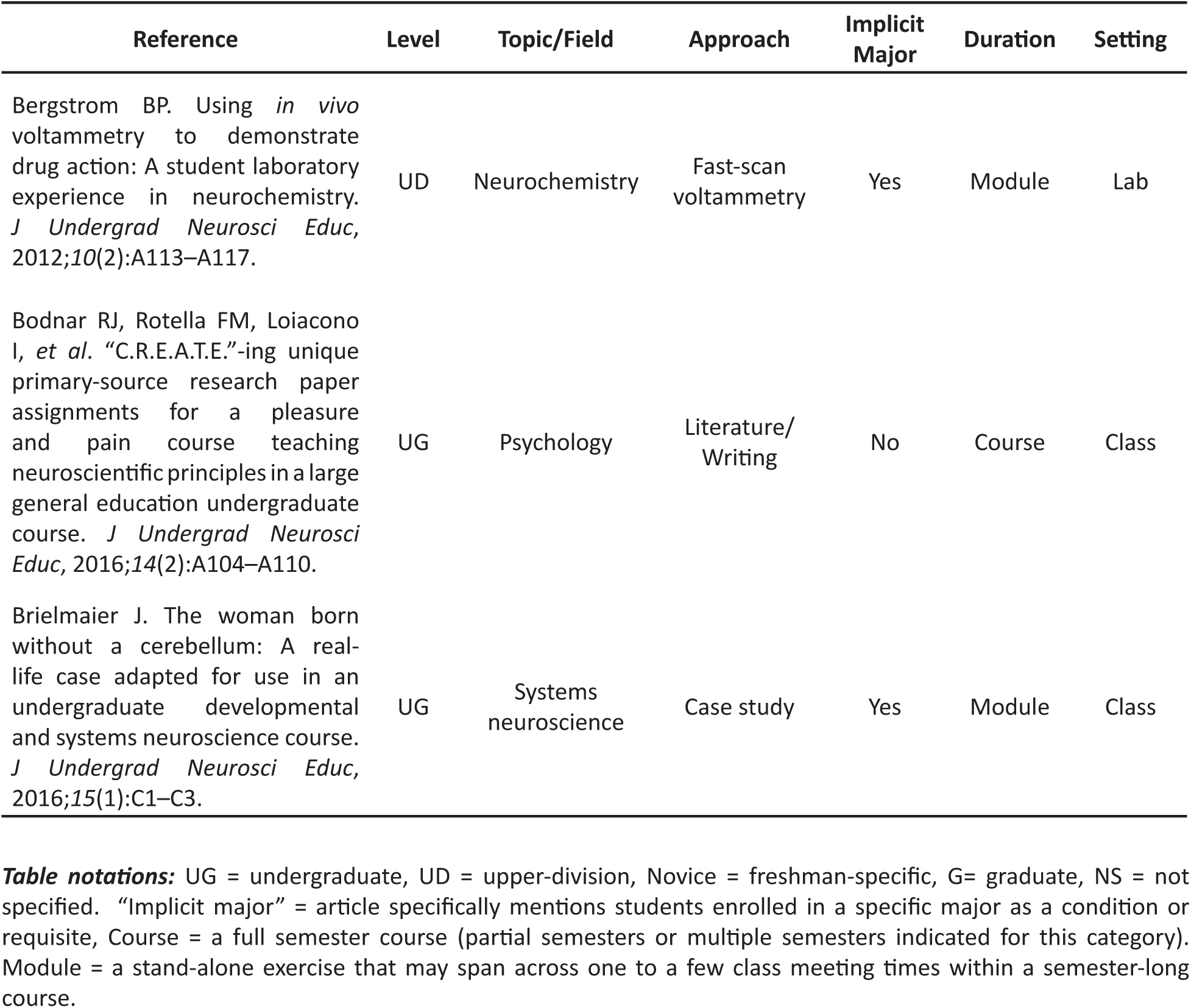
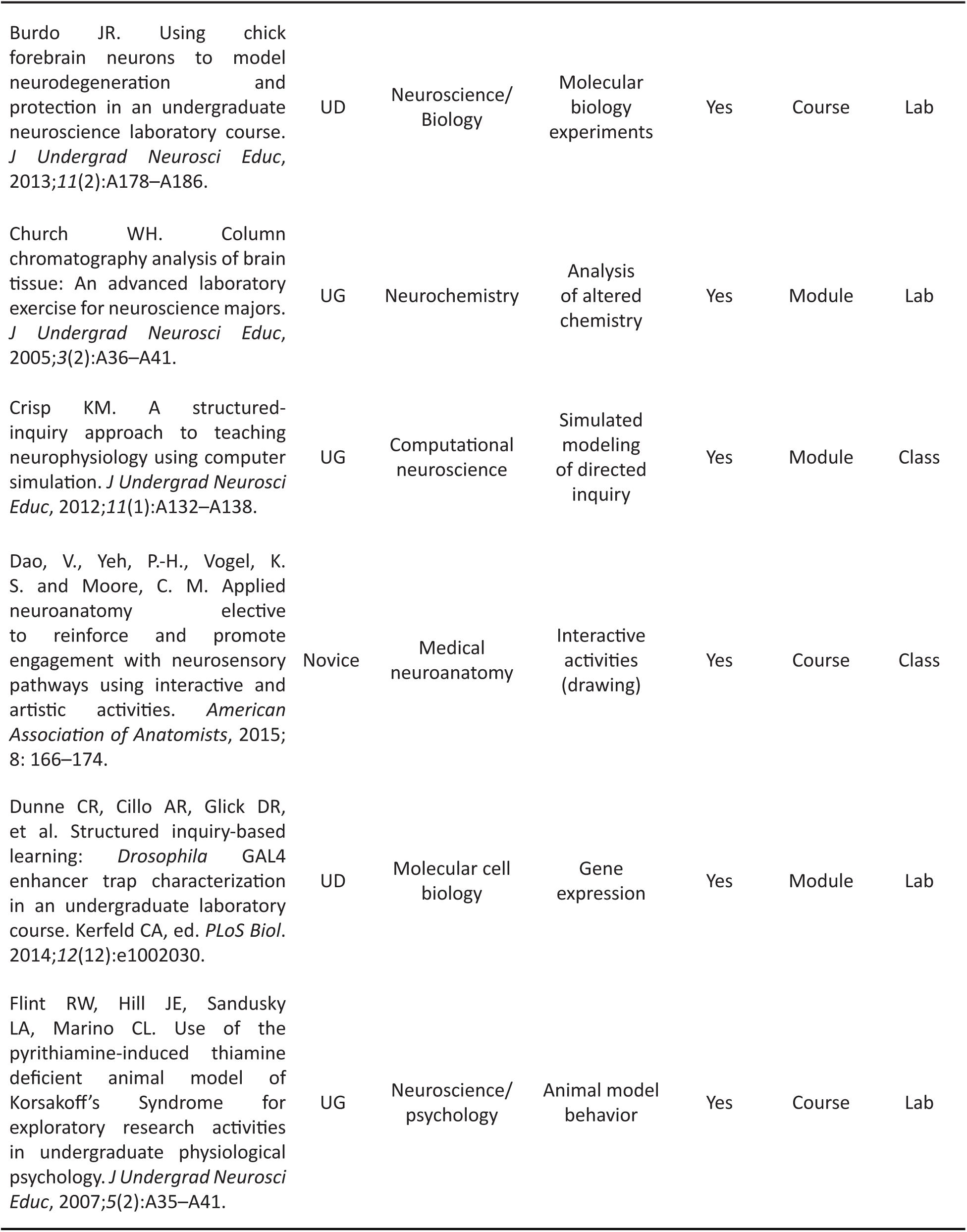
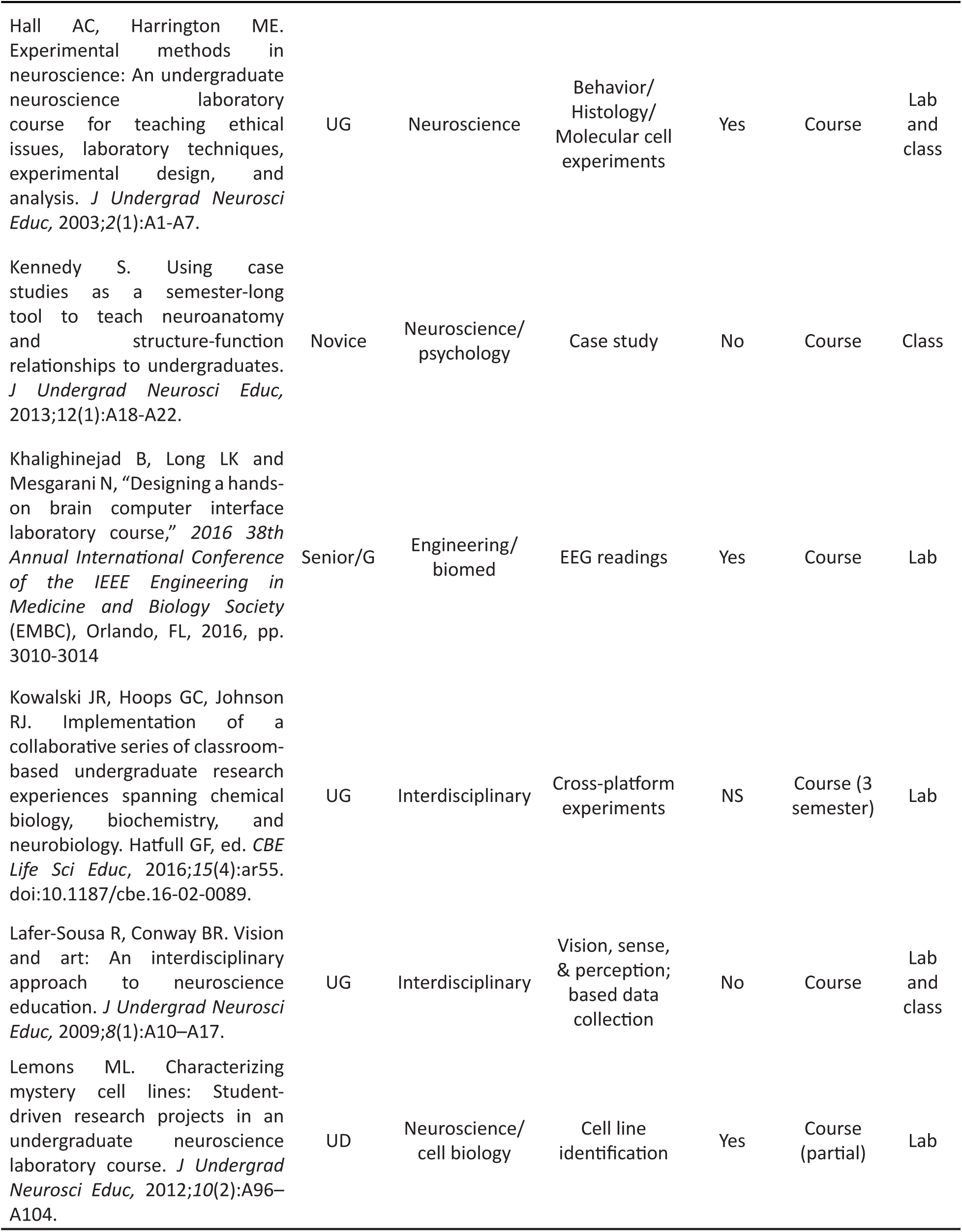
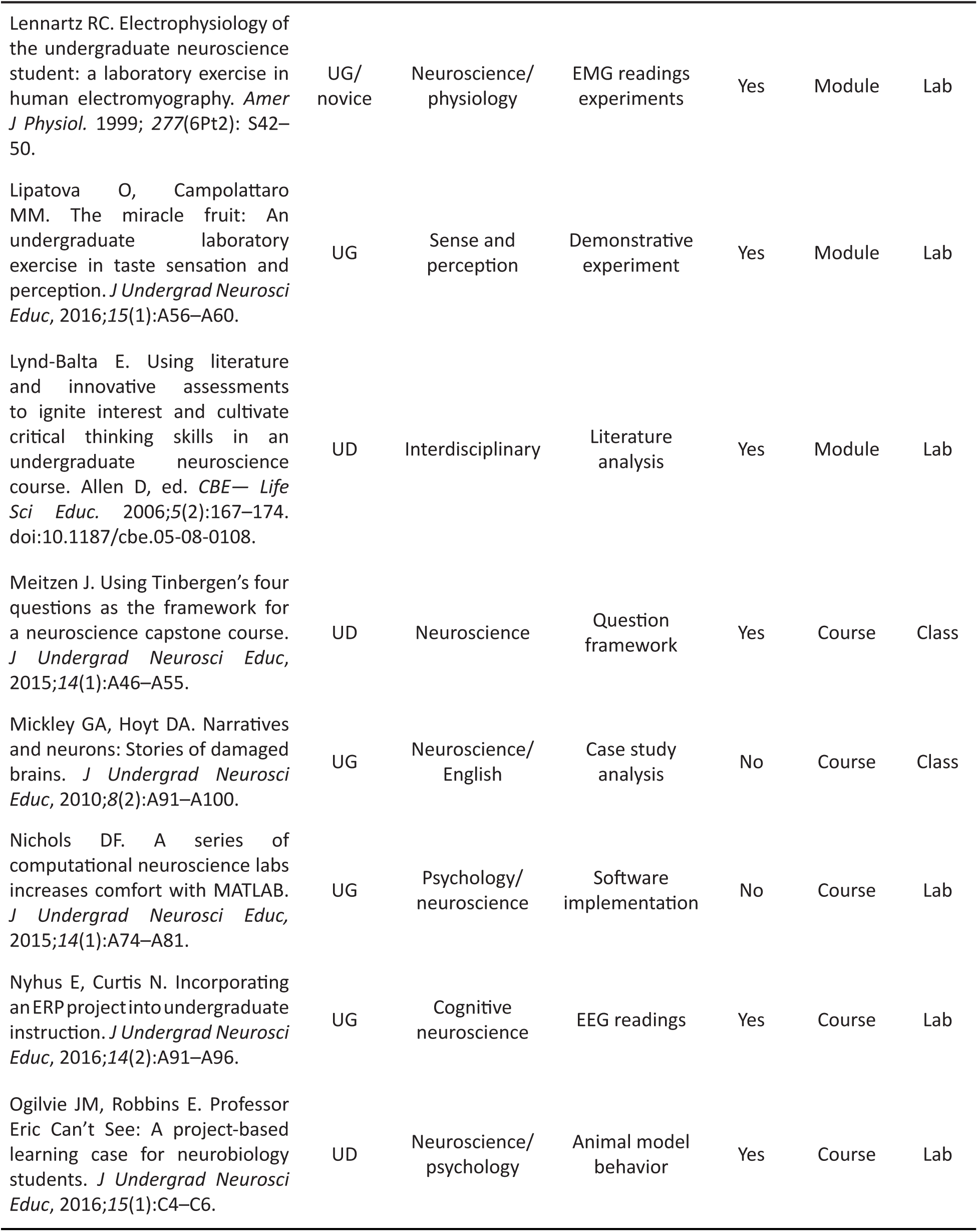
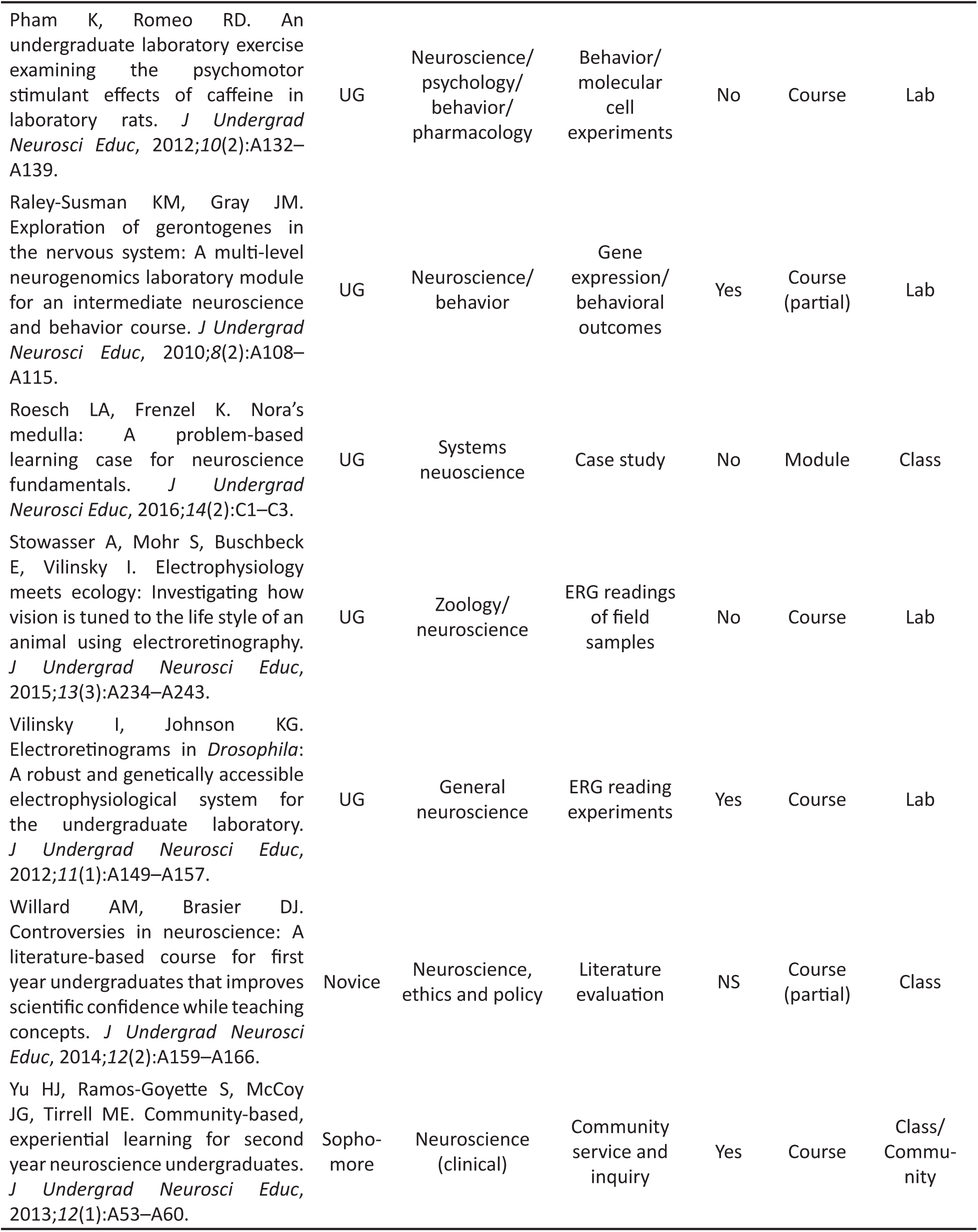
Published reports of curricula and/or implementations of neuroscience CUREs.

## D’Arcy *et al*. Supplemental Materials II: Scoring and Rubric for Mapping Neuromapping Scoring Rubric (NSR): Development and Implementation

### Purpose

Traditional summative evaluation approaches(*e.g.*, examsorlaboratory reports summarizing experiments) often inadequately assess application and integration of skills by evaluating an end result only. This practice ignores the iterative nature of the CURE/authentic research environment wherein students areafforded multipleopportunitiestocorrectandimprovetheir work. In developing the NSR we present a tool for formative evaluation of student products that assists instructors in identifying students’ functional understanding or misconceptions, in providing directed feedback to students, and in streamlining the vetting process for the final cytoarchitectural maps. To promote standardized evaluation and quality control of aggregate student data over the course of multiple student cohorts, categories of the NSR (an example of a completely scored NSR is presented in **Table SM2.1**) address the students’ ability 1) to recognize the difference between the model (atlas templates) and the individual subject examined, 2) to accurately identify the anatomical structures while acknowledging and accommodating variability introduced by differences among animals or by errors in the plane of section, and 3) to provide a representational translocation of experimental data to a standard reference atlas (*see* **Companion Illustration to Table SM2.1**). This rubric does not evaluate the histological skills of the student; *i.e.*, whether their methods produced tissue damage, tissue distortion, tissue folds and wrinkles, bubbles in mounting media, or variation in stain intensity and image noise.

### Development

Three reviewers with varying degrees of expertise in brain mapping (novice, experienced in non-hypothalamic regions, experienced in hypothalamic regions) independently scored 18 sample maps of variable quality. Poor initial interrater reliability scores among specific categories were corrected by clearly defining scoring criteria and reducing scale resolution to a binary score, as presented below. For data presented in this study, two reviewers scored student maps independently (k = 0.881; p < 0.001; 95% CI (0.650, 1.112)).

### Neuromapping Scoring Rubric: Scoring Guidelines

*Note*: Student products should meet the following categories with >70% compliance to receive a score of “1.” Students falling below the desired threshold will be awarded a score of “0.” All results should be discussed with students in order to provide them with formative opportunities to revise their maps. All scores of “1” should be reviewed with students to ensure that proper rationale (where applicable) has been ascribed. Collectively, this tool is to be used for assessing student progress and comprehension as well as for facilitating product improvement for the purpose of enhancing scientific merit. This is not to be used toward a course grade.

#### 1. System and Logic: Knowledge and Application **(Figs. SM2.1, SM2.2**)

##### Consistency in demarcation throughout

**Table SM2.1.** Scoring rubric for atlas-based neuroanatomical mapping.

|  |  | Group 1 | Group 2 | Group 3 | Group 4 | M (SD) |
| --- | --- | --- | --- | --- | --- | --- |
| System and Logic | Consistency in demarcation throughout | 1 | 1 | 1 | 1 | 1.00 (±0.0) |
|  | File layers organized accessibly | 1 | 1 | 1 | 1 | 1.00 (±0.0) |
|  | File layers labeled informatively | 1 | 1 | 1 | 1 | 1.00 (±0.0) |
|  | File layer groups organized and labeled logically | 1 | 1 | 1 | 1 | 1.00 (±0.0) |
| Atlas Proficiency | Dominant atlas level correctly identified | 1 | 1 | 1 | 1 | 1.00 (±0.0) |
|  | Correct atlas level assignment for plane shift | 1 | 1 | 1<sup>a</sup> | 0 | 0.75 (±0.5) |
|  | Recognition of a shift in plane of section | 1 | 1 | 1<sup>b</sup> | 1 | 1.00 (±0.0) |
|  | Indicated the axis of change in plane | 1 | 1 | 1 | 0 | 0.75 (±0.5) |
| Anatomical Proficiency | Major grey matter regions Identified | 1 | 1 | 1 | 1 | 1.00 (±0.0) |
|  | Major grey matter regions correctly bounded | 1 | 1 | 1 | 1 | 1.00 (±0.0) |
|  | Sub-regions identified | 1 | 1 | 1 | 0 | 0.75 (±0.5) |
|  | Sub-regions correctly bounded | 0 | 1 | 1 | 0 | 0.50 (±0.6) |
| Signal Identification | Complete cataloging of immunopositive cell bodies | 1 | 0 | 1 | 1 | 0.75 (±0.5) |
|  | Complete cataloging of immunopositive fibers | 1 | 1 | 1 | 1 | 1.00 (±0.0) |
|  | Correct signal representation/placement | 1 | 0 | 1 | 0 | 0.50 (±0.6) |
|  | Interaction and localization zones indicated | 0 | 0 | 0 | 0 | 0.00 (±0.0) |
| Map Score |  | 14 | 13 | 15 | 10 | 13 (±2.16) |

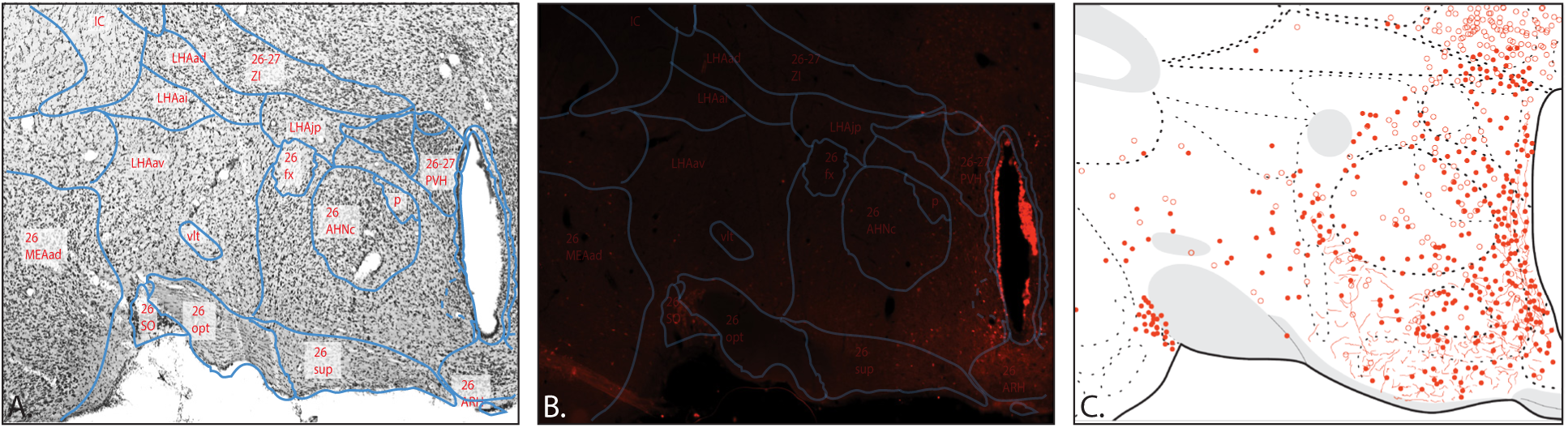
Companion Illustration to Table SM2.1. A. Sample images were obtained from Group 3 of Cohort 2 to illustrate student parcellation efforts of Nissl-stained tissue (×10 magnification). Clear labels are used throughout, and atlas and anatomical proficiency are evident in the boundaries drawn for gray and white matter regions. **B.** Immunofluorescence image of calbindin immunoreactivity from Group 3 files provide an example of student mounting and staining capabilities. Students aligned the image of the Nissl-stained section and its parcellation to the immunofluorescent single-channel image, thus providing regional boundaries for the immunofluorescent section. The student indicated the atlas level(s) and region names within the delineated image. Note that mounting artifacts alter the registration of the parcellated-boundaries overlay with the main landmarks (white matter tracts, ventricle) in the underlying image. **C.** Signal identification and representation for the data for calbindin-immunoreactive cells is presented for Level 26. Note, in this example, students adopted the convention of using an open circle to denote cells that expressed calbindin immunoreactivity with clearance from the zone of the nucleus in lieu of using that notation to indicate weakly immunoreactive cells. The underlying atlas template is from Swanson (2004) (available at https://larrywswanson.com) and is reproduced and modified here under the conditions of a Creative Commons BY-NC 4.0 license (https://creativecommons.org/licenses/bync/4.0/legalcode).

Demarcation of parcellation boundaries, fibers, and cells must maintain a consistent line width, line style convention (example: solid for confirmed boundaries, dotted for ambiguous), color setting (RGB or CMYK value is consistent), and circle size (including outline) (**Fig. SM2.2**).

##### File layers organized accessibly

File layers should be ordered to allow for consistent viewing of pertinent data (**Fig. SM2.1**).

##### File layers labeled informatively

Names of file layers must accurately describe the information contained in the layer (example: Nissl Parcellation) (**Fig. SM2.1**).

##### File layer groups organized and labeled logically

Representations of immunopositive staining should be organized by antigen and should be grouped with corresponding immunohistochemical (IHC) images. Example: Each IHC antigen should have its own parent layer with associated sublayers for cells and fibers to allow rapid toggling of information (**Fig. SM2.1**).

**Figure SM2.1:**
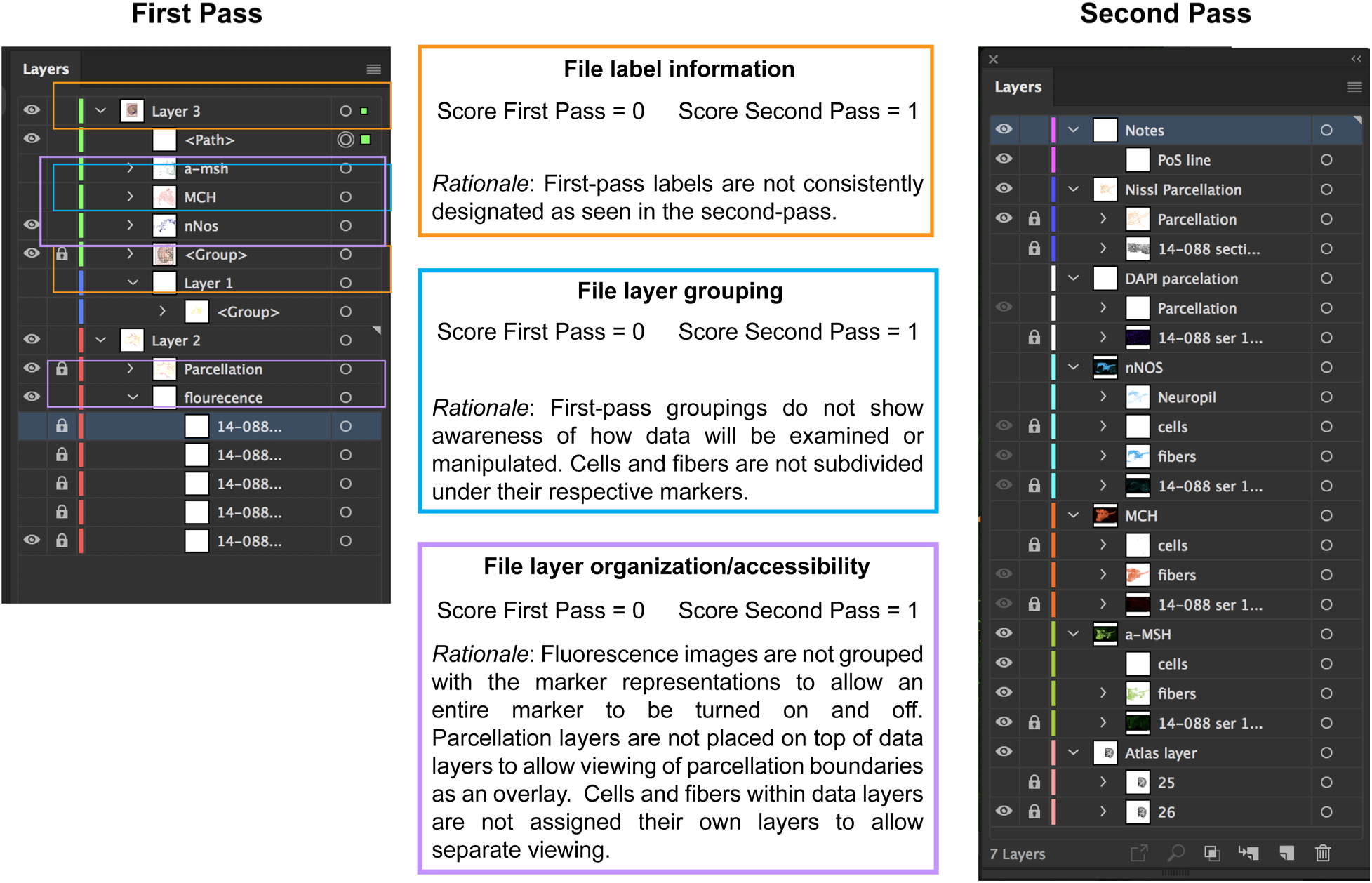
Scoring examples for the *System and Logic* category. Adobe Illustrator allows image files and objects (type, and vector paths). to be organized into data layers. These layers can be controlled independently (including locking or viewing), assigned descriptive titles, and assigned additional properties. As an example of the latter, the color bar depicted on the left can be switched to easily denote the pseudocolor that was used to represent a given IHC antigen (*e.g.*, nNOS, MCH, and αMSH shown here from Cohort 1). In these screen captures of the data layer organization, first-pass efforts betray a less sophisticated understanding of the use of the software and/or the practical manipulation of the data layers than do second-pass efforts. Note that screen shots of First- and Second-Pass *Layers* dialog boxes are student-generated and preserve any errors they have made.

#### 2. Atlas proficiency: Knowledge, application, and comprehension **(Fig. SM2.2**)

##### Dominant atlas level correctly identified

The atlas level that most closely represents the analyzed image should be identified in the working map (often a file naming convention). If the transition between planes is such that two layers are equally represented through the region of interest, the two atlas levels should have clear indications of their regions of applicability (**Fig. SM2.2**). The final maps for an atlas level should represent only the data that apply to that level.

##### Recognition of a shift in plane of section

Distinguishing shifts in the shape of gray matter regions across atlas levels should be denoted either by assigning the corresponding atlas level to each identified gray matter region/sub-region or by using marking conventions, such as a grid or line indicating the transition between atlas levels (**Fig. SM2.2**).

**Figure SM2.2:**
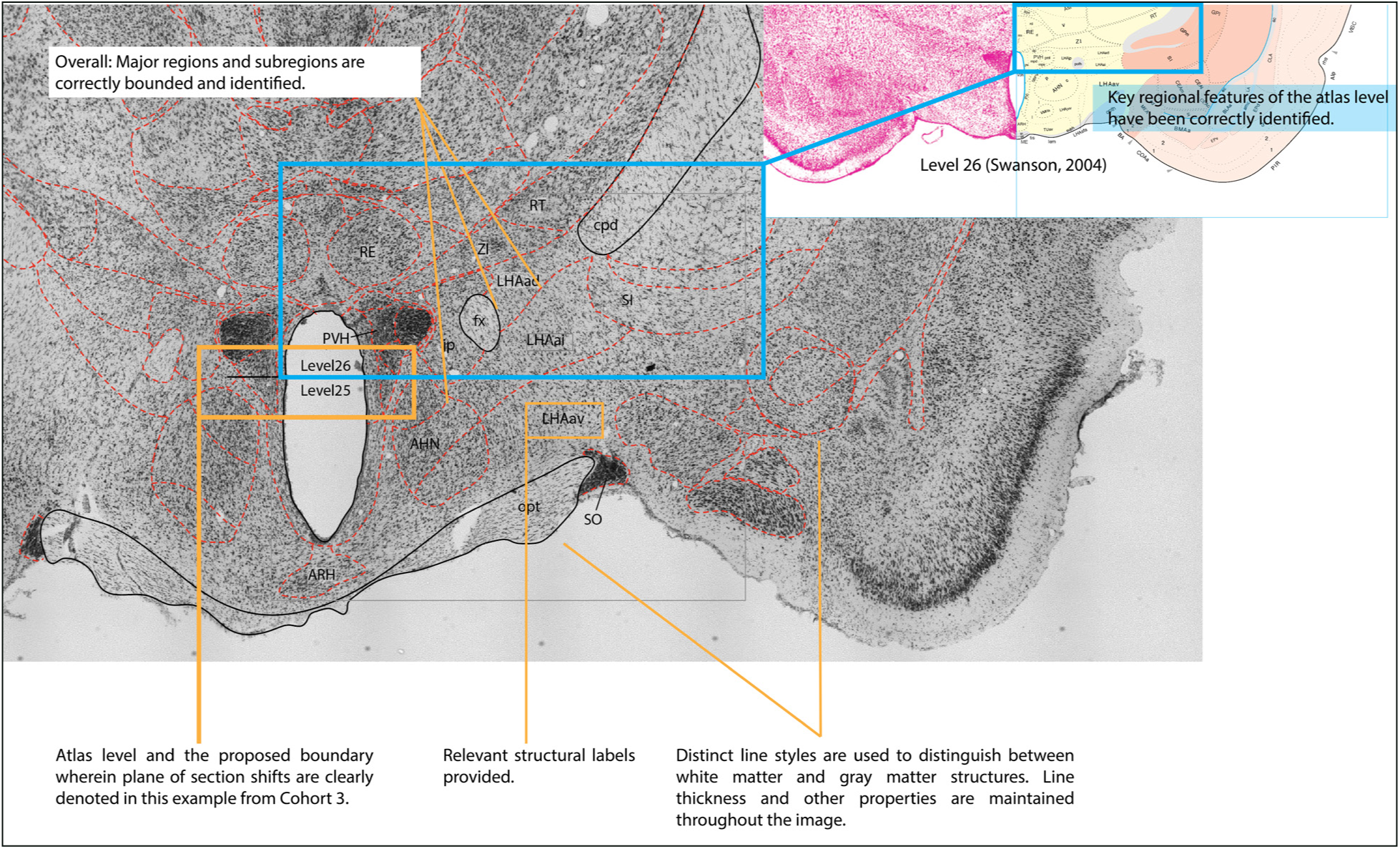
Plane of section analysis in a parcellated image of a Nissl-stained section. This example, drawn from Cohort 3, provides an example of good consistency in demarcation (*System and Logic* category) and evidence of clear atlas and anatomical proficiency in all categories. Note the students’ ability to distinguish boundaries based on cytoarchitectural distinctions established in the image of the Nissl-stained section (×10 magnification) rather than arbitrarily attempting to overlay the representational atlas boundaries directly. A line (partially obscured by the blue frame) has been provided to indicate where the shift in atlas level takes place and the approximate direction in which plane-of-section shift occurs. Atlas levels have been correctly assigned based on key anatomical features (an example of an atlas template for the level is provided and the region of interest is framed in *blue*). Key structures have been identified (region of interest = LHA, ZI), including white matter tracts used as fiducials. The atlas level in the inset is from Swanson (2004) (available at https://larrywswanson.com) and is reproduced and modified here under the conditions of a Creative Commons BY-NC 4.0 license (https://creativecommons.org/licenses/bync/4.0/legalcode).

##### Correct atlas level assignment for plane shift

Identified atlas levels should correctly correspond to the reference atlas based on key structures or other fiducials.

##### Indicated the axis of change in plane

Though the axis (direction across which the plane of section transitions between atlas levels) may be inferred, in most cases, by assessing level labels per structure in the atlas notes, additional notation of the directional trend in (and point in which) the plane of section traverses atlas levels is recommended. Alternatively, the change of plane may be indicated by a line or grid depending upon the complexity of the tissue distortion observed.

#### 3. Anatomical proficiency: Knowledge, application, comprehension, analysis (**Figs. SM2.2, SM2.3**)

##### Major gray matter regions identified

All hypothalamic gray matter regions (list provided by instructor) should be identified and parcellated if present in the image.

##### Major gray matter regions correctly bounded

Parcellations of major structures, such as the paraventricular hypothalamic nucleus (PVH), should have an accompanying rationale to support boundaries.

##### Sub-regions identified

Structures of interest that have internal subdivisions should include all clearly definable substructures (for example, for the LHA at Level 26, the structures include the LHAad, LHAai, LHAav, LHAjp).

##### Sub-regions correctly bounded

Refer to recommendations for major gray matter regions.

#### 4. Signal identification (**Figs. SM2.3, SM2.4**)

##### Complete cataloging of immunopositive cell bodies

After working with the course instructor to establish thresholds for what constitutes an immunopositive cell, students should clearly indicate all immunopositive somata.

##### Complete cataloging of immunopositive fibers

Scoring parameters for this criterion are consistent with those established for cataloging immunopositive somata.

##### Correct signal representation/placement

Data that are transferred to the reference map should reflect the appropriate localization and orientation within the structure and maintain the positional relationship with surrounding structures.

##### Interaction and localization zones indicated

Regions indicating signal trends (consistency with which immunoreactive cells and

**Figure SM2.3:**
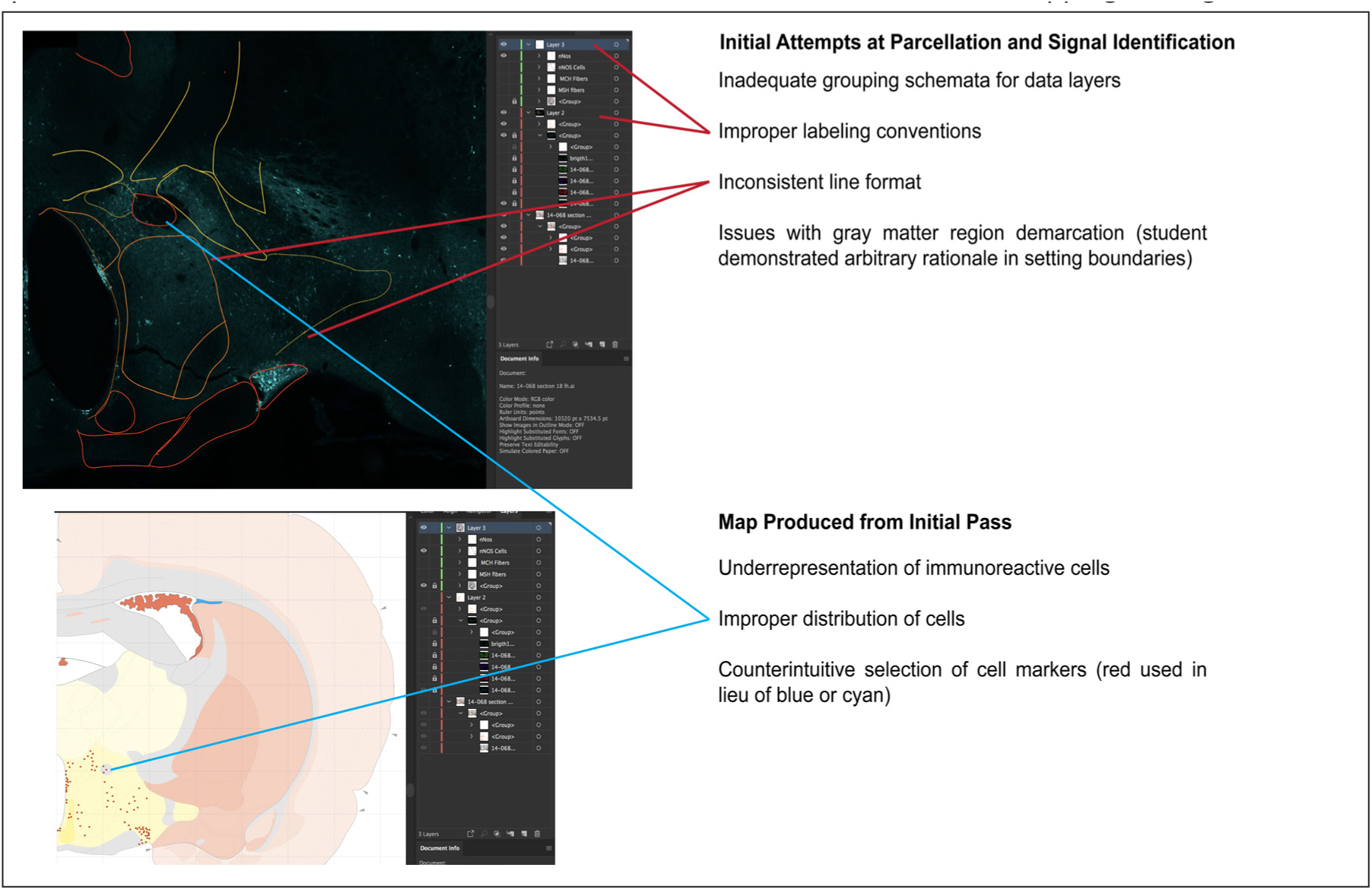
Preliminary attempts of mapping prior to rubric-guided feedback. This example, drawn from Cohort 1, shows a preliminary attempt at parcellation and mapping for nNOS-immunoreactive cell bodies. Note that several discrepancies exist not only in data layer organization, but also in the consistency and accuracy of boundary placement. Note, too, the underrepresentation of cell bodies and the inaccurate migration of data onto the atlas map as most clearly evident for cells were drawn within the fornix in the atlas template. The atlas map is from Swanson (2004) (available at https://larrywswanson.com) and is reproduced and modified here under the conditions of a Creative Commons BY-NC 4.0 license (https://creativecommons.org/licenses/bync/4.0/legalcode).

**Figure SM2.4:**
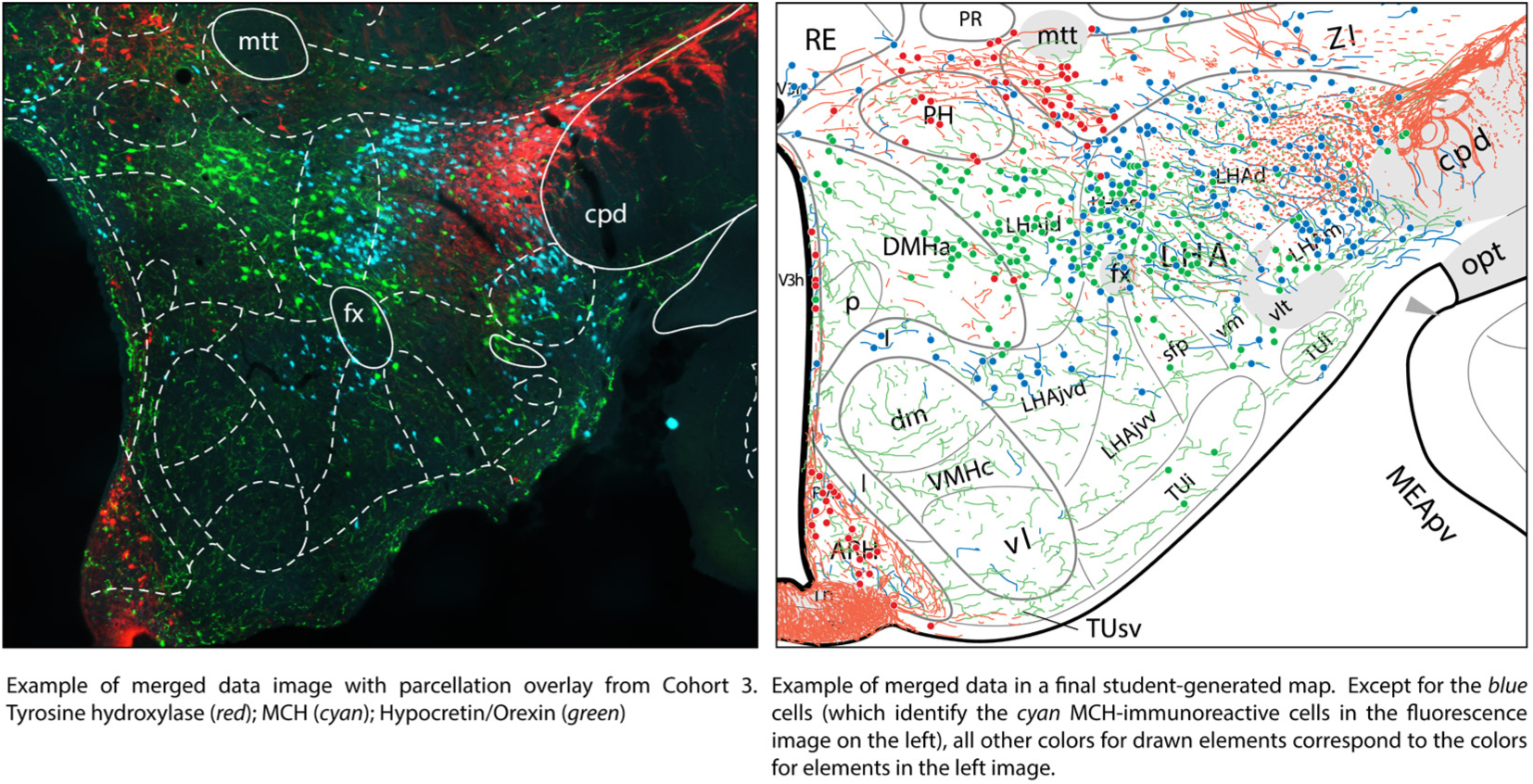
Example of final data representation on an atlas level in comparison to the original data image. These images (reproduced from **Fig. 6** in the main text) illustrate that the distribution of cell somata and fibers are translated to the final map in a manner that preserves the positional relationships of signal within the depicted structures. The image at the left was taken at ×10 magnification. The atlas level is from Swanson (2004) (available at https://larrywswanson.com) and is reproduced and modified here under the conditions of a Creative Commons BY-NC 4.0 license (https://creativecommons.org/licenses/bync/4.0/legalcode).

## D’Arcy *et al*. Supplemental Materials III: Syllabus for Brain Mapping & Connectomics

### Brain Mapping and Connectomics Syllabus

#### Contact information

##### Scheduled lab hours

###### Lab goal

The goal of the HHMI Brain Mapping lab is to generate real-world research that is published in a peer-reviewed journal. This makes the Brain Mapping lab unique – you will not just learn a variety of research techniques, and you will not just mimic real research by participating in a semester-long project. You will participate in an actual dissertation project, and your work will be shared with the scientific community. Along the way, you will pick up histological and anatomical research techniques, expand your critical thinking skills, be exposed to professional science writing, and be asked to produce written work in a professional format. In order to achieve the primary course goal of becoming an author on a peer-reviewed scientific publication, you must: be prepared to put in a large amount of time working on the project, and produce high-quality work that meets the stringent standards of the scientific journal this project will be published in.

###### Project goal

You will be working on the Khan lab’s hypothalamic chemoarchitecture project. The goal of this project is to map the distribution patterns of neurons expressing specific chemicals in the rat hypothalamus and to analyze the interactions among these neurons. The hypothalamus is a region of the brain that is famous for participating in control of homeostasis, autonomic and neuroendocrine output, and some motivated behaviors. Disorders of the hypothalamus can potentially affect sleep patterns, metabolic and cardiovascular health, sexual development, and temperament, among other things. There are many different neuronal circuits present in the hypothalamus, but where they are and what form they take is largely unknown, which makes it impossible to understand how the hypothalamus carries out its functions or how it might be disordered in certain medical conditions. Because different neurons express different neurotransmitters, one way to see brain circuits is as a pattern of interacting chemical systems. The hypothalamic chemoarchitecture (chemical architecture) project is concerned with describing, in a very precise and rigorous way, various chemical systems present in the hypothalamus as well as their interactions, as a way of gaining information about hypothalamic circuitry. The project findings will be mapped to a reference atlas (the Swanson 2004 atlas of the adult male rat brain), so that they will be easy for other investigators to use and to integrate with their own data.

Because there are so many different neurotransmitters expressed in the hypothalamus, this is a long-term project which may easily take decades to complete. It will be published as a series of papers that each focus on a particular set of neurotransmitters that we will work together to select as a class.

###### Student contribution to the project

In groups of four students, you will stain rat brain tissue to visualize functional markers in the brain and map their distribution patterns to the Swanson atlas. Each group will be responsible for staining and mapping one brain. Each group member will be responsible for an equal amount of this work. Some of the mapping will be completed as a group; whatever is left will be divided equally amongst the group members. You will prepare a report of your findings and present your maps to your instructor for quality assessment and inclusion in a paper to be published in a peer-reviewed journal, assuming that quality requirements are met.

##### Policies

###### Protection of data

Because you will be conducting actual research, the information you generate in this lab is the intellectual property of [institution omitted]. It does not belong to you. All staining procedures must be written up in your lab notebook, which cannot be removed from the laboratory. You will be allowed to remove the Illustrator files containing your maps from the lab, so that you may work on them at home or at the library, but an up-to-date copy of mapping files must remain on the lab computers at all times. If questions arise amongst the scientific community concerning the staining, your lab notebooks will be referred to. If questions arise as to the integrity of the mapping, your Illustrator files will be referred to. Your lab notebook is considered a legal document. This is serious business.

###### Required items

1. Pen
2. Sharpie
3. Number 0 round (or number 0 liner) watercolor brush with natural bristles. The brush tips should not have hairs that splay out to the side and should come to a fine point. Avoid artificial bristles like golden taklon or white nylon. These brushes will be used in manipulating slices of tissue-paper-thin brain, so choose your tool carefully!
4. Lab coat

###### Excused absences, unexcused absences, and tardiness

Absences may be excused for the following reasons: professional development (such as attendance at a conference), military duties, membership on a [institution omitted] sports team that requires you to leave town for a game, hospitalization or illness, or death in the family. If you know you will need to miss a day of lab, inform your instructor as soon as possible. The instructor must be presented with supporting documentation for any absence to be considered excused, regardless of the reason for the absence. Having an excused absence does not excuse you from completing any assignments that may have been due on that date. Additionally, coordinate with your team members in order to figure out division of duties, etc., so that your absence does not impose undue burden to your teammates.

One unexcused absence will result in the reduction of your grade by a letter-grade, and two or more unexcused absences will result in an automatic grade of F.

If you are delayed from arriving to class on time and able to safely contact your team members, notify them of what is going on and work with them to resolve any work imbalance. This lab reflects real life as a scientist, and, thus, requires real-life problem solving with your research team.

###### Academic dishonesty

In accordance with the policies of [institution omitted], academic dishonesty is considered wholly intolerable. Academic dishonesty is essentially any form of cheating. Students found to commit academic dishonesty will be disciplined following [institution omitted] established procedures, which you can read about here: [link omitted].

###### Plagiarism

Plagiarism is a form of academic dishonesty and will not be tolerated whatsoever. If any part of an assignment submitted by a student is plagiarized, the entire assignment will be considered plagiarized and will count as a zero (0). Particularly, flagrant and deliberate plagiarizers will be referred to the Dean of Students, in accordance with [institution omitted] policies on academic dishonesty.

###### ADA accessibility

Students with disabilities who require accommodations must contact the Center for Accommodations and Support Services (CASS) as soon as possible at [link omitted]. The CASS office is [location omitted], and their website is [link omitted].

###### Use of HHMI brain mapping lab equipment

This lab is funded by a grant from the HHMI. Your tuition and lab fees do not contribute to the setup and maintenance of this facility or its equipment. Therefore, your use of the facility and equipment is a privilege, not a right. You may not be present in the lab without the course instructor being present, and you may not use any equipment without the permission of your course instructor. If you violate these stipulations or use the facilities and/ or equipment in a disrespectful or irresponsible manner, the TA or course instructor may throw you out of lab and disallow you from attending lab.

##### Grading

|  |  |
| --- | --- |
| Participation and Attendance | 40% |
| Notebook Check/Quizzes | 20% |
| Project Development and Abstract | 20% |
| Blogging | 10% |
| Final Maps* | 10% |

##### Assignments

###### Participation and attendance

This is a research lab. We are finding the answers to questions no one else has answered before. As such, coming into the lab is essential. Your team, your class, and your collaborating scientist are depending on you to do your part.

###### Notebook check/quizzes

As a scientist, there is nothing more valuable than keeping clear, detailed, and honest records of what you did and how you did it in order to produce the data you observe, record, and report. Keeping accurate records will be essential, and one of the ways we check notebook content is to conduct notebook-based in-class quizzes. You will have the opportunity to review your notebook contents at the start of class in preparation for the quiz/potentially use your notebook to furnish information during the quiz.

###### Blogging

Taking time to think about your science is all a part of the process. Blogging will be similar to your lab notebook in that you will document your progress each week, including any staining that you did and any mapping that you completed. How many sections did you mount? How many sections did you parcellate, etc.? But it is also where you can reflect on problems you encounter, articles you have read, and how your data fits into a bigger picture.

###### Project development and abstract

Your teams will work together to develop a project to tackle. Part of this will involve reading through scientific papers, and part of it will involve a careful selection of what you will test and how you will set up the experiment (including what areas of the brain we want to examine, why they are important to examine, and what are some signals we want to examine and why). Additionally, we will work together to develop individual, group, and class abstracts that summarize your findings and determine how to present your data on a poster for a professional conference. Participation and demonstration of critical thinking are keys to succeeding in this grade category.

###### Final maps

For the grade, your instructor will evaluate not the final map, but each step taken to obtain it. Throughout the semester, you will be observed in how well you contribute to your team’s maps, your intellectual discussions that take place during the course of mapping, your care in maintaining ordered and clearly-labeled layers, your attention to detail in maintaining line thickness and color, your honesty in noting regions of ambiguity, and your rationale behind your brain region boundaries (parcellations). *It is not about the destination, but the journey.

While not assigned a grade, your final maps will be subject to the critique and examination of the scientific community. Your Illustrator files will be used to assess the quality of your maps by the lab Principle Investigator. High-quality maps will be those that adhere to the reference standards established, that reflect uncertainties in parcellations and transfers carefully, that are assigned to an atlas level(s) correctly and in a thoughtful manner (meaning you didn’t just draw a line across the section), and that demonstrate a subtle and refined understanding of the chemoarchitecture underlying parcellations. Such maps will contribute to the poster and future publications. If your maps meet these standards for inclusion, you become eligible for inclusion as an author in subsequent professional communications and reports using this data (see your instructor for further details).

##### Team Contract

I will be on time.

I will be respectful of my teammates and their time.

I will respect the safety of others and will obey all safety rules and encourage my teammates to do so.

When working as a group, I will contribute intellectually to the task at hand.

I will do my part to complete research outside of lab that my teammates are depending on me for. I will not allow my teammates to wallow in egregious amounts of my unfinished work.

If I fail in these duties to my team, I understand that my final grade will suffer a loss of points in an amount corresponding to the severity of my failure of my team.

***__________________________***

***Signature***

***__________________________***

***Printed Name***

***__________________________***

***Date***

## References

American Association for the Advancement of Science (2011) Vision and change in undergraduate biology education: A call to action. Washington, D.C.: American Association for the Advancement of Science.

Auchincloss LC, Laursen SL, Branchaw JL, Eagan K, Graham M, Hanauer DI, Lawrie G, McLinn CM, Pelaez N, Rowland S, Towns M, Trautmann NM, Varma-Nelson P, Weston TJ, & Dolan EL (2014) Assessment of course-based undergraduate research experiences: A meeting report. CBE Life Sci. Educ. 13, 29–40.

Badalà F, Nouri-Mahdavi K & Raoof DA (2013) Making a difference in science education: the impact of undergraduate research programs. Am. Educ. Res. J. 50(4), 683–713.

Ballen CJ, Blum JE, Brownell S, Hebert S, Hewlett J, Klein JR, McDonald EA, Monti DL, Nold SC, Slemmons KE, Soneral PAG, & Cotner S (2017) A call to develop course-based undergraduate research experiences (CUREs) for nonmajors courses. CBE Life Sci. Educ. 16(2), mr2.

Bonwell CC, & Eison JA (1991) Active Learning: Creating Excitement in the Classroom. ASHE-ERIC Higher Education Report No. 1. Washington, DC: The George Washington University, School of Education and Human Development.

Brown RA, & Swanson LW (2013) Neural Systems Language: A formal modeling language for the systematic description, unambiguous communication, and automated digital curation of neural connectivity. J. Comp. Neurol. 521, 2889–2906.

Buonaccorsi VP, Boyle MD, Grove D, Praul C, Sakk E, Stuart A, Tobin T, Hosler J, Carney SL, Engle MJ, Overton BE, Newman JD, Pizzorno M, Powell JR, & Trun N (2011) GCAT-SEEKquence: Genome Consortium for Active Teaching of Undergraduates through increased faculty access to next-generation sequencing data. CBE Life Sci. Educ. 10, 342–345.

Buonaccorsi V, Peterson M, Lamendella G, Newman J, Trun N, Tobin T, Aguilar A, Hunt A, Praul C, Grove D, Roney J, & Roberts W (2014) Vision and change through the Genome Consortium for Active Teaching Using Next-Generation Sequencing (GCAT-SEEK). CBE Life Sci. Educ. 13, 1–2.

Burnett KAS, Pinales BE, Perez EJ, Rodarte D, Cardona AM, Galvan KJ, Hernandez GG, Lezama AC, Lorenzana KT, Vasquez A, Parada P, Paz JI, Rascon J, Thomason R, Bautsta K, Barnes J, D’Arcy C, Khan AM (2018) High spatial resolution mapping of anorexigenic neuropeptides expressed in the hypothalamus: A chemoarchitecture study in the adult male rat. Program No. 680.24. 2018 Neuroscience Meeting Planner. San Diego, CA: Society for Neuroscience, 2018. Online.

Call GB, et al. (2007) Genomewide clonal analysis of lethal mutations in the *Drosophila melanogaster*eye: Comparison of the X chromosome and autosomes. Genetics 177, 689–697.

Chen J, et al. (2005) Discovery-based science education: Functional genomic dissection in Drosophila by undergraduate researchers. PLoS Biol. 3(2), e59.

Chickering AW, & Gamson ZF (1987) Seven principles for good practice in undergraduate education. A.A.H.E. Bull. pp. 3–6; March.

Corwin LA, Runyon C, Robinson A, & Dolan EL (2015) The laboratory course assessment survey: a tool to measure three dimensions of research-course design. CBE Life Sci. Educ. 14, 1–11.

D’Arcy CE, Khan AM & Olimpo JT (2016a) Implementation of a connectomics course-based undergraduate research experience in introductory biology. 13th International Sun Conference on Teaching and Learning, 17–18 Mar, El Paso, TX.

D’Arcy C, Martinez AM, Wells CE, Khan AM & Olimpo JT (2016b) Implementation of a brain mapping course-based undergraduate research experience in introductory biology: Impacts on novices’ competency and affect. Program No. 22.06SA. 2016 Neuroscience Meeting Planner. San Diego, CA. Society for Neuroscience. Online.

D’Arcy C, Martinez A, Aranda LF, Cervantes HFL, Chacon LE, Cordero RP, Fernandez V, Garcia GA, Holguin S, Jacquez A, Miramontes TG, Montaño B, Muñoz PC, Valenzuela IR, Yu JS & Khan AM (2016c) 6Elaboration of hypothalamic chemoarchitecture of the adult male rat: A high spatial resolution mapping study of melanin-concentrating hormone, hypocretin/orexin, and calbindin immunoreactivities in multiple subjects. Program No. 453.07. 2016 Neuroscience Meeting Planner. San Diego, CA. Society for Neuroscience. Online.

Della Mea V, Maddalena E, Mizzaro S, Machin P, & Beltrami CA (2014) Preliminary results from a crowdsourcing experiment in immunohistochemistry. Diagn. Pathol. 9(Suppl 1), S6.

Dolan, EL (2017) Undergraduate research as curriculum. Biochem. Mol. Biol. Educ. 45(4), 293–298.

Elgin SCR, Hauser C, Holzen TM, Jones C, Kleinschmit A, & Leatherman J (2016) The GEP: Crowd-sourcing big data analysis with undergraduates. Trends Genet. 33(2), 81–85.

Estrada M, Wookcock A, Hernandez PR, & Schultz PW (2011) Toward a model of social influence that explains minority student integration into the scientific community. J. Educ. Psychol. 103(1), 206–222.

Feynman RP (1985) “Surely You’re Joking Mr. Feynman!” Adventures of a Curious Character. New York: Bantam Books.

Flores-Robles G, Negishi K, Pacheco RA, Enriquez A, Acevedo E, Avila B, Dominguez E, Hernandez EE, Medina A, Mejia E, Novoa MA, Provencio AT, Renteria FD, Sifuentes E, Tellez Y, & Khan AM (2017) Hypothalamic chemoarchitecture of the adult male rat: Further elaboration of results from a high spatial resolution longitudinal mapping study. Program No. 604.03. 2017 Neuroscience Meeting Planner. Washington, DC. Society for Neuroscience, 2017. Online.

Frantz KJ, Demetrikopoulos MK, Britner SL, Carruth LL, Williams BA, Pecore JL, DeHaan RL, & Goode CT (2017) A comparison of internal dispositions and career trajectories after collaborative versus apprenticed research experiences for undergraduates. CBE Life Sci. Educ. 16(1), ar1.

Geerling JC, & Loewy AD (2006) Aldosterone-sensitive neurons in the nucleus of the solitary tract: efferent projections. J. Comp. Neurol. 497, 223–250.

Hahn J (2010) Comparison of melanin-concentrating hormone and hypocretin/orexin peptide expression patterns in a current parceling scheme of the lateral hypothalamic zone. Neurosci. Lett. 468, 12–17.

Hanauer DI, & Hatfull G (2015) Measuring networking as an outcome variable in undergraduate research experiences. CBE Life Sci. Educ. 14, 1–10.

Handelsman J, Ebert-May D, Beichner R, Bruns P, Chang A, DeHaan R, Gentile J, Lauffer S, Stewart J, Tilghman SM, & Wood WB (2004) Scientific teaching. Science 304, 521–522.

Herrick CJ (1926) Brains of rats and men: A survey of the origin and biological significance of the cerebral cortex. Chicago: University of Chicago Press. Available online at the Internet Archive: http://www.archive.org/details/brainsofratsandm031896mbp. Accessed on June 9, 2017.

Irshad H, Oh EY, Schmolze D, Quintana LM, Collins L, Tamimi RM, & Beck AH (2017) Crowdsourcing scoring of immunohistochemistry images: Evaluating performance of the crowd and an automated computational method. Sci. Rep. 7, 43286.

Jeffery E, Nomme K, Deane T, Pollock C, & Birol G (2016) Investigating the role of an inquiry-based biology lab course on student attitudes and views toward science. CBE Life Sci. Educ. 15(4), ar61.

Kerman IA, Bernard R, Rosenthal D, Beals J, Akil H, & Watson SJ (2007) Distinct populations of presympathetic-premotor neurons express orexin or melanin-concentrating hormone in the rat lateral hypothalamus. J. Comp. Neurol. 505, 586–601.

Khan AM (2013) Controlling feeding behavior by chemical or gene-directed targeting in the brain: what’s so spatial about our methods? Front. Neurosci. 7, 182.

Khan AM, Grant AH, Martinez A, Burns GAPC, Thatcher BS, Anekonda VT, Thompson BW, Roberts ZS, Moralejo DH, & Blevins JE (2018a) Mapping molecular datasets back to the brain regions they are extracted from: Remembering the native countries of hypothalamic expatriates and refugees. Advances in Neurobiology, 21,101–193.

Khan AM, Perez J, Wells CE, & Fuentes O (2018b) Computer vision evidence supporting craniometric alignment of rat brain atlases to streamline expert-guided, first-order migration of hypothalamic spatial datasets related to behavioral control. Frontiers in Systems Neuroscience, 12(Article 7), 1–29.

Klein A, & Hirsch J (2005) Mindboggle: A scatterbrained approach to automate brain labeling. NeuroImage 24, 261–280.

Kowalski JR, Hoops GC, & Johnson RJ (2016) Implementation of a collaborative series of classroom-based undergraduate research experiences spanning chemical biology, biochemistry, and neurobiology. CBE Life Sci. Educ. 5, ar55.

Martinez A, Barraza Escudero LM, Castro D, Chavez SA, Coronado M, Gallegos S, Pineda Sanchez A, Ruiz MSP, Ruiz VG, Negishi K, & Khan AM (2018) Hypothalamic chemoarchitecture of the adult male rat: High spatial resolution mapping of copeptin, LIM homeobox 6, and melanin-concentrating hormone. Program No. 680.23. 2018 Neuroscience Meeting Planner. San Diego, CA: Society for Neuroscience, 2018. Online.

Martinez A, Navarro VI, Arias K, Barnett J, Carreon A, Castaneda B, Flores M, Magadan J, Heredia D, Hooper J, Lopez T, Lozano A, Mendez S, Mercer N, Munoz A, Rosario Mojica L, Sanchez R, Sierra K, Sotelo D, Stevens X, Toccoli A, Verma Y, Vizcarra H, & Khan AM. (2019) The Hypothalamic Chemoarchitecture Project, Year 5: High-spatial resolution atlas-based mapping of neuronal populations expressing melanin concentrating hormone, hypocretin/orexin, and calbindin in the hypothalamus of the adult male rat. Submitted to the Society for Neuroscience Annual Meeting (to be held in Chicago, IL; Oct 19–23, 2019).

National Research Council (1996) From analysis to action: Undergraduate education in science, mathematics, engineering, and technology. Washington, D.C.: National Academies Press.

National Research Council (2003) BIO2010: Transforming undergraduate education for future research biologists. Washington, D.C.: National Academies Press.

OED Online. Oxford University Press. (March, 2017). http://0-www.oed.com.lib.utep.edu/view/Entry/376403?redirectedFrom=crowdsourcing& (accessed June 04, 2017).

Oleson D (2011) Crowdsourcing scientific research: Leveraging the crowd for scientific discovery. Available online at: https://www.crowdflower.com/scientific-research/. (Accessed on June 4, 2017).

Olimpo JT, Fisher GR, & DeChenne-Peters SE (2016) Development and evaluation of the Tigriopus course-based undergraduate research experience: Impacts on students’ content knowledge, attitudes, and motivation in a majors introductory biology course. CBE Life Sci. Educ. 15, ar72.

Owens MT, & Tanner KD (2017) Teaching as brain changing: Exploring connections between neuroscience and innovative teaching. CBE Life Sci. Educ. 16(fe2), 1–9.

Pineda Sanchez A#, Enriquez A#, K Negishi, Santarelli AJ, Khan AM*, Poulos AM.* (2019). Atlas-based mapping of Fos-immunoreactive neurons in the medial prefrontal cortex and ventral hippocampus in association with context fear in the juvenile and adult male rat. Accepted for the Society for Neuroscience Annual Meeting (to be held in Chicago, IL; Oct 19–23, 2019). # co-first authors; *co-senior authors.

Rodenbusch SE, Hernandez PR, Simmons SL & Doln EL (2016) Early engagement in course-based research increases graduation rates and completion of science, engineering, and mathematics degrees. CBE Life Sci. Educ. 15, 1–10.

Roskams J, & Popovic Z (2016) Power to the people: Addressing Big Data challenges in neuroscience by creating a new cadre of citizen neuroscientists. Neuron 92, 658–664.

Santarelli AJ, Khan AM, & Poulos AM (2018) Contextual fear retrieval-induced Fos expression across early development in the rat: An analysis using established nervous system nomenclature ontology. Neurobiol. Learn. Mem. 155, 42–49.

Simmons DM, & Swanson LW (2009) Comparing histological data from different brains: Sources of error and strategies for minimizing them. Brain Res. Rev. 60, 349–367.

Swanson LW (2004) Brain Maps: Structure of the Rat Brain, 3rd ed. Elsevier, Amsterdam.

Swanson LW, Sanchez-Watts G, & Watts AG (2005) Comparison of melanin-concentrating hormone and hypocretin/orexin mRNA expression patterns in a new parceling scheme of the lateral hypothalamic zone. Neurosci. Lett. 387, 80–84.

Wells CE, Acosta A, Aldrete D, Carrion A, Castro L, Escapita S, Espinoza E, De La Fuente K, Garrett A, Gomez N, Gomez N, Hernandez-Casner C, Luevano M, Lopez A, Martinez D, Mendoza E, Ortega M, Perez M, Rangel E, Reza E, Rivera J, Roman C, Rosas A, Seade-Galindo C, Teran J, Unpingco J, Valdez S & Khan AM (2015a) Next generation interaction maps of hypothalamic circuits controlling survival behaviors. Workshop: Hypothalamic Circuits for Control of Survival Behaviors; 2015 Sep 27; Janelia Farm Research Campus, Ashburn, Virginia.

Wells CE, Acosta A, Aldrete D, Carrion A, Castro L, Escapita AE, Espinoza E, De La Fuente K, Garrett A, Gomez A, Gomez N, Hernandez-Casner C, Luevano M, Lopez A, Martinez D, Mendoza E, Ortega M, Perez M, Rangel E, Reza E, Rivera J, Roman C, Rosas A, Seade-Galindo C, Teran J, Unpingco J, Valdez S & Khan AM (2015b) Hypothalamic chemoarchitecture in the adult male rat: Creating canonical atlas maps for co-visualized immunoreactive peptidergic neuronal populations (α-MSH, nNOS, MCH) and their fiber systems in multiple brains. Program No. 616.08. 2015 Neuroscience Meeting Planner. Chicago, IL. Society for Neuroscience. Online.

Weston TJ, & Laursen SL (2015) The Undergraduate Research Student Self-Assessment (URSSA): Validation for use in program evaluation. Cell Biol. Educ. 14, ar33–ar33.

Yao ST, Gourand S, Paton JFR, & Murphy D (2005) Water deprivation increases the expression of neuronal nitric oxide synthase (nNOS) but not orexin-A in the lateral hypothalamic area of the rat. J. Comp. Neurol. 490, 180–193.

Zséli G, Vida B, Martinez A, Lechan RM, Khan AM, & Fekete C (2016) Elucidation of the anatomy of a satiety network: Focus on connectivity of the parabrachial nucleus in the adult rat. J. Comp. Neurol. 524, 2803–2827.

